# Local ancestry inference identifies robust evidence of selection in Neolithic Europe

**DOI:** 10.64898/2026.04.23.720248

**Authors:** Georgia Mies, Iain Mathieson

## Abstract

During the European Neolithic, migrating Anatolian farmers admixed with local hunter-gatherers, coinciding with major shifts in diet, environment, and lifestyle that imposed strong selective pressures. Local ancestry inference is widely used to detect selection following admixture, but most methods were developed and validated on present-day populations. Their performance in ancient DNA – where reference panels are smaller, data are sparser, and admixture is more ancient – remains unresolved. We benchmark eight local ancestry inference methods on 176 imputed Neolithic genomes. While individual-level ancestry estimates are highly correlated across methods, inferred tract lengths and admixture time estimates vary by an order of magnitude. Overall, we recommend Gnomix or RFMix for general use. We also investigated our ability to detect natural selection using LAI. Integrating results across methods and replicating across methods and in two independent datasets (n=378 and 1,121) we identify a robust ancestry deviation at *FADS1/2*, consistent with adaptation on metabolism. We also identify *IRAK4* (innate immunity) as a candidate locus, but with less consistent signal across methods. Finally, we replicate previous reports of excess hunter-gatherer ancestry at the HLA, but these results are inconsistent across methods and suggest that they may be affected by bias in local ancestry inference. Our findings demonstrate that while local ancestry inference recovers biologically meaningful signals in ancient genomes, results can be sensitive to the methods used for inference, particularly in complex regions like the HLA. Method choice critically influences inferred ancestry patterns and selection signals, underscoring the importance of multi-method validation.

## Introduction

Early farmers from the Anatolian peninsula initiated the European agricultural transition (Neolithic; 10-7 thousand years ago) by introducing domesticated animals, agricultural subsistence, and related lifeways through both migration and subsequent admixture with local European hunter-gatherer groups (Brandt et al. 2013; Lipson et al. 2017). The resulting admixed Neolithic population is an example of a two-way admixture event accompanied by substantial environmental and cultural change (Lipson et al. 2017; Ju and Mathieson 2021; Childebayeva et al. 2022). Genomic patterns of admixture from this period therefore reflect both demographic history and adaptation to novel environments (Korunes and Goldberg 2021; Cuadros-Espinoza et al. 2022). Farmer ancestry makes up approximately 80% of admixed Neolithic genomes, with hunter-gatherer ancestry accounting for the remaining 20% (Haak et al. 2015; Allentoft et al. 2024), increasing over time as regional hunter-gatherer groups were absorbed into farming populations (Lipson et al. 2017; Rivollat et al. 2020).

Admixture is ubiquitous across human populations and contributes to shaping genetic variation, introducing ancestry blocks that may harbor adaptive variants (Korunes and Goldberg 2021). Selection acting on these linked ancestry blocks produces local signatures of selection, observed as excess ancestry from one source population relative to the genome-wide background. Studying post-admixture adaptation sheds light on how populations adapted to new environments, diets, pathogens, and lifestyles, especially during major transitions like the Neolithic. During the agricultural transition, natural selection driven by strong environmental and cultural shifts may have acted on variation introduced by either farmers adapted to agricultural lifestyles, or hunter-gatherers adapted to the European environment. Identifying variants that increased in frequency after the onset of agriculture can provide insight into traits that were biologically important during this period.

Local ancestry inference (LAI) – the classification of ancestry along the genome – provides a framework for studying fine-scale admixture patterns, allowing for estimation of admixture timing from ancestry tract lengths, and identification of adaptive admixture. LAI has been widely applied to present-day admixed populations, but its application to ancient DNA remains limited. Ancient DNA presents several challenges for LAI: low coverage, genotype uncertainty, smaller and unbalanced reference panels (Davy et al. 2023), post-mortem damage, and greater temporal distance between source and admixed populations. Most LAI methods were developed and validated using large, high-coverage present-day datasets with well-matched reference panels (Oliveira et al. 2024). Their performance under the conditions typical of ancient DNA, remains poorly characterized. Biases in imputation or phasing of ancient DNA could also bias LAI accuracy, distorting estimates of ancestry proportions, tract lengths, admixture timing, and signals of selection. Nonetheless, several papers have used LAI in ancient individuals to make inference about selection and disease risk. Analysis of ancestry deviation in the admixed Neolithic found elevated hunter-gatherer ancestry at the HLA, consistent with adaptation to local pathogens (Davy et al. 2023; da Silva et al. 2025), and elevated farmer ancestry at the pigmentation gene *SLC24A5*, associated with lighter skin pigmentation (Ju and Mathieson 2021; Davy et al. 2023). LAI of Bronze Age and present-day individuals has been used to identify associations between ancestry at the HLA and multiple sclerosis (Barrie et al. 2024) and between ancient ancestry and disease risk in present-day populations (Marnetto et al. 2022; Pankratov et al. 2024).

We benchmarked eight widely used LAI methods using 176 imputed and phased genomes from admixed Neolithic individuals (Allentoft et al. 2024). We compared inferred ancestry proportions, tract length distributions, and signatures of selection across methods and replicated selection signals in two independent datasets (n=378 and 1,121). Finally, we show that by systematically comparing multiple LAI approaches in a well-characterized ancient admixture scenario we can distinguish robust signals of adaptive admixture from methodological artifacts.

## Results

### Local ancestry inference is sensitive to sample size, posterior filtering, and imputation

We evaluated the performance of LAI methods on 176 admixed Neolithic individuals with published imputed genomes ranging from 1,450 to 8,676 years old (mean: 5,542 years) and spanning Western and Central Europe (Figure 1; Gamba et al. 2014; Lazaridis et al. 2014; Olalde et al. 2014; Skoglund et al. 2014; Allentoft et al. 2015; Günther et al. 2015; Jones et al. 2015; Olalde et al. 2015; Cassidy et al. 2016; Kılınç et al. 2016; González-Fortes et al. 2017; Martiniano et al. 2017; Fernandes et al. 2018; Valdiosera et al. 2018; Antonio et al. 2019; Brace et al. 2019; González-Fortes et al. 2019; Malmström et al. 2019; Sánchez-Quinto et al. 2019; Schroeder et al. 2019; Brunel et al. 2020; Scorrano et al. 2022; Allentoft et al. 2024). For each LAI method, source populations consisted of 48 Western hunter-gatherers and 7 Anatolian farmers, hereafter referred to as hunter-gatherers and farmers. Prior to LAI, we filtered autosomal SNPs for overlap between imputed data and the 1240k SNP set from the Allen Ancient DNA Resource (AADR; Mallick et al. 2024), applying minor allele frequency (MAF) > 0.01 and imputation quality > 0.8 filters, yielding 572,223 autosomal SNPs. For the X chromosome, 160,150 SNPs remained after filtering for MAF > 0.05 in both the imputed dataset and the 1000 Genomes Project (Auton et al. 2015) reference panel.

**Figure 1.**
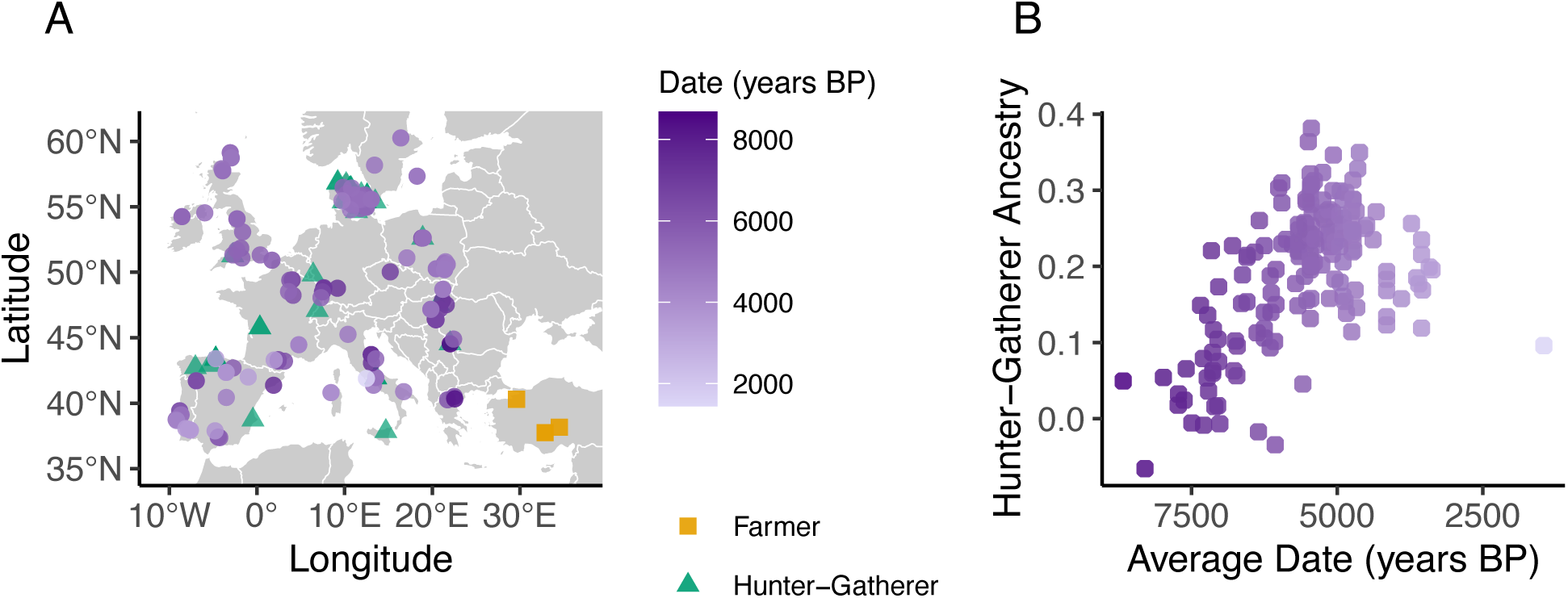
Geographic and temporal distributions of admixed Neolithic individuals. **A)** Geographic locations of individual collection sites. **B)** Hunter-gatherer global ancestry proportions estimated by qpAdm for each admixed individual, plotted by estimated date (years before present). Point color indicates the date of the individual.

To benchmark LAI performance on these ancient genomes, we selected widely used LAI methods representing distinct algorithmic classes, including six that we ran ourselves: Gnomix (Hilmarsson et al. 2026), RFMix V1.9 (Maples et al. 2013), Recomb-Mix (Wei et al. 2025), Mosaic (Salter-Townshend and Myers 2019), simpLAI (Oliveira et al. 2024), Ancestry HMM (Corbett-Detig and Nielsen 2017), and published results from a seventh method AncestralPaths (Pearson and Durbin, unpublished data; Allentoft et al. 2024; Table 1). We were unable to use Flare (Browning et al. 2023) since it reported zero hunter-gatherer ancestry for all individuals. Flare is specifically designed for larger sample sizes, so it is perhaps unsurprising that it is not useable on this small sample. AncestralPaths reported inflated hunter-gatherer ancestry estimates on chromosomes 6 and 9 relative to other chromosomes (Figure S1), which we corrected by adjusting each chromosome’s ancestry proportion to match the genome-wide average. In the absence of known ground truth for ancient Neolithic genomes, we assess LAI methods by comparison to allele frequency-based estimates, known temporal constraints on admixture timing, and replication across independent datasets.

**Table 1.** Local ancestry inference methods. Summary of the local ancestry inference methods evaluated, listing the primary inference algorithm and whether the method includes ancestry-switch correction (rephasing) to account for phasing errors in the input data. For consistency, we additionally applied post hoc ancestry-switch correction to the output of all methods before downstream analyses.

| Method | Publication | LAI algorithm | Rephasing |
| --- | --- | --- | --- |
| Gnomix | Hilmarsson et al. 2026 | Deep learning classifier (CNN + XGBoost) with smoothing | Yes |
| RFMix V1.9 | (Maples et al. 2013) | Random forest classifier with CRF/HMM smoothing | Yes |
| Recomb-Mix | Wei et al. 2025 | Hidden Markov model (HMM) with graph optimization | Yes |
| Mosaic | Salter-Townshend and Myers 2019 | Hidden Markov Model (HMM) | Yes |
| simpLAI | Oliveira et al. 2024 | Haplotype matching (minimum mismatch; 7 HG 7 farmer references) | No |
| Ancestry HMM | Corbett-Detig and Nielsen 2017 | Hidden Markov model (HMM) using read counts | No |
| AncestralPaths | Pearson and Durbin, unpublished data; Allentoft et al. 2024 | Neural network | No |

To assess overall estimation accuracy, we compared genome-wide ancestry proportions from LAI methods against the allele frequency-based approaches qpAdm (Haak et al. 2015) and ADMIXTURE (Alexander et al. 2009). qpAdm estimated an average hunter-gatherer ancestry of 0.20 (range: −0.10 to 0.41; Figure 1), while unsupervised ADMIXTURE (K = 2) reported an average of 0.16 (range: 0 to 0.39). All LAI methods overestimated individual hunter-gatherer ancestry relative to qpAdm when all reference individuals were used, suggesting varying degrees of bias toward the overrepresented hunter-gatherer source population (48 hunter-gatherers vs. 7 farmers, compared to roughly 1:5 ancestry proportions; Figure S2). Gnomix showed substantially lower bias (mean difference 0.04), whereas all other methods differed from qpAdm by 0.21-0.52. To test whether this bias was due to sample size or the nature of the data, we ran LAI on African American (ASW) admixed individuals from the 1000 Genomes Project (Auton et al. 2015) with similarly unbalanced reference panels. RFMix, Gnomix and Mosaic produced unbiased results, while Recomb-Mix, simpLAI, and Ancestry HMM were biased (Figure S3).

This suggests that these three methods are fundamentally sensitive to sample size, while RFMix and Mosaic are sensitive only under some conditions, perhaps when source populations are relatively closely related. Among the evaluated methods, Gnomix produces the least biased ancestry estimates in both ancient and present-day datasets.

To mitigate the bias across methods, we optimized source sample sizes for each LAI method by subsampling hunter-gatherers, except for AncestralPaths where we retained published calls using the original sample sizes. Using either 3 or 7 hunter-gatherers alongside 7 farmers yielded the best results (Figures 2 and S2). Additionally, filtering LAI calls by posterior probability > 0.9 improved concordance with qpAdm for methods that report posteriors (Figure S2). Individual hunter-gatherer ancestry proportions from all methods were correlated with qpAdm estimates, though correlation strength and bias varied (Figure 2).

**Figure 2.**
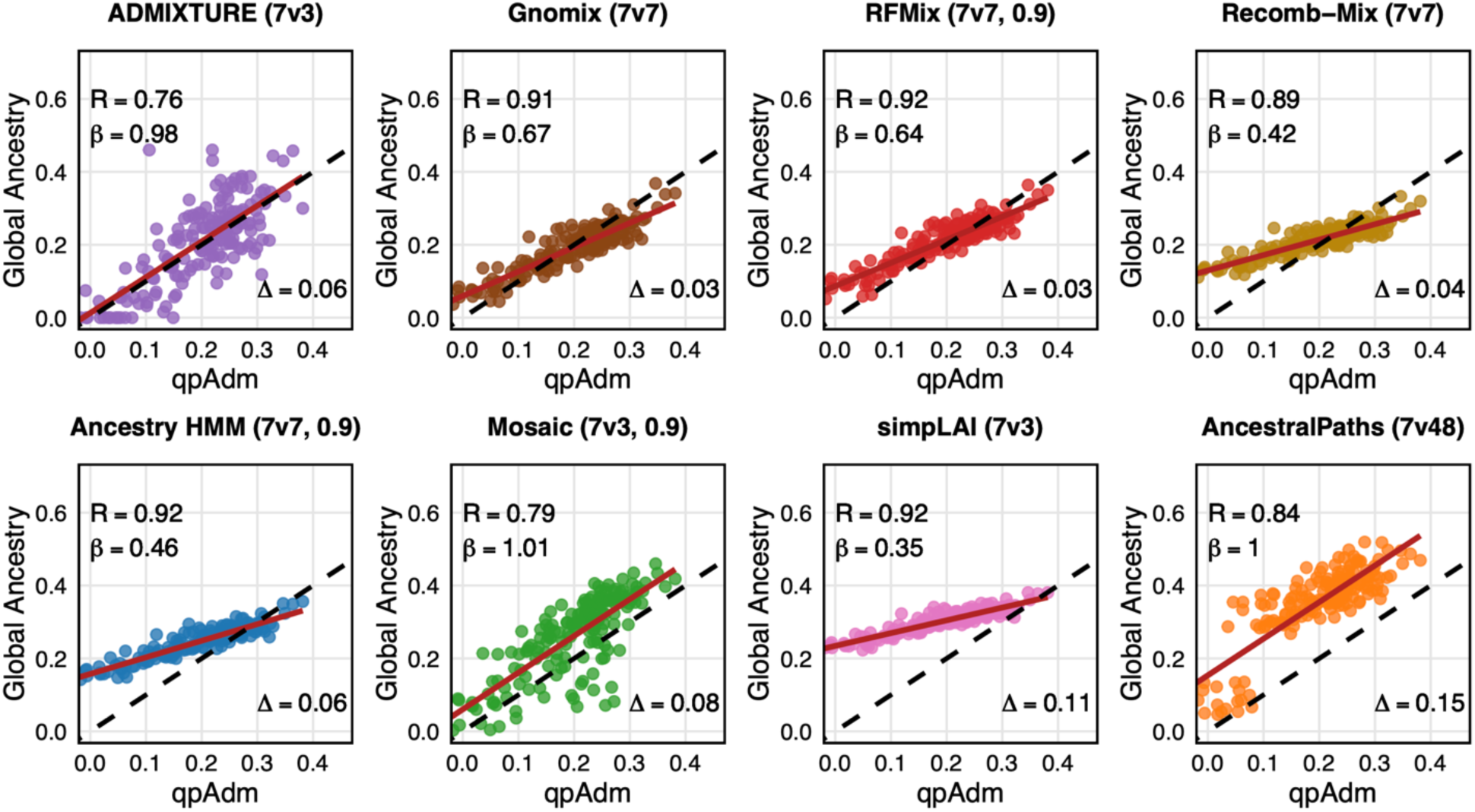
Whole-genome ancestry estimates by method. Correlations between global ancestry proportions estimated by ADMIXTURE and LAI methods versus qpAdm. Each point represents one of the 176 admixed Neolithic individuals. R denotes the Pearson correlation between the method and qpAdm individual estimates. β indicates the slope of the regression line between the method and qpAdm estimates. Δ represents the average absolute difference in global ancestry between the LAI method and qpAdm. The dashed line corresponds to y = x for qpAdm estimates, while the red line shows the best-fit linear regression between qpAdm and the LAI method. Sample sizes used for the best-fit regressions are shown in parentheses with the method title, indicating 7 farmers versus N hunter-gatherers (7vN) and whether method calls were filtered by a 0.9 posterior probability threshold. Methods are ordered by lowest to highest mean squared error from qpAdm estimates.

Next, we investigated biases related to data processing. Global ancestry proportions estimated from imputed versus unimputed data were highly similar for allele frequency-based methods (qpAdm and ADMIXTURE), indicating that imputation in itself does not bias ancestry proportions (Figure S4). On the other hand, Ancestry HMM – the only LAI method that can run on unimputed data – does show a substantial difference in global ancestry between imputed and unimputed input suggesting that LAI methods in general may be sensitive (Figure S4). LAI of the X chromosome revealed no differences in ancestry proportion or tract lengths between males (with perfect phasing) and females (phased as autosomes), indicating that phasing errors do not substantially impact LAI results (Figures S5 and S6).

### Local ancestry methods disagree about the location and length of local ancestry tracts

Although optimizing source sample sizes and applying posterior probability filtering improved concordance of global ancestry estimates between LAI methods, substantial disagreement remained in local ancestry. We evaluated local ancestry concordance by comparing genome-wide correlations of hunter-gatherer ancestry proportions in 100 kb bins between pairs of methods.

Pearson correlations ranged from 0.08 to 0.5 with AncestralPaths consistently showing the lowest correlations relative to other methods (Figure 3). Across individuals, genome-wide local ancestry correlations ranged from 0.15 to 0.43 (mean = 0.32; 100 kb bins; Figure S7), indicating substantial variation in the inferred genomic distribution of ancestry among methods.

**Figure 3.**
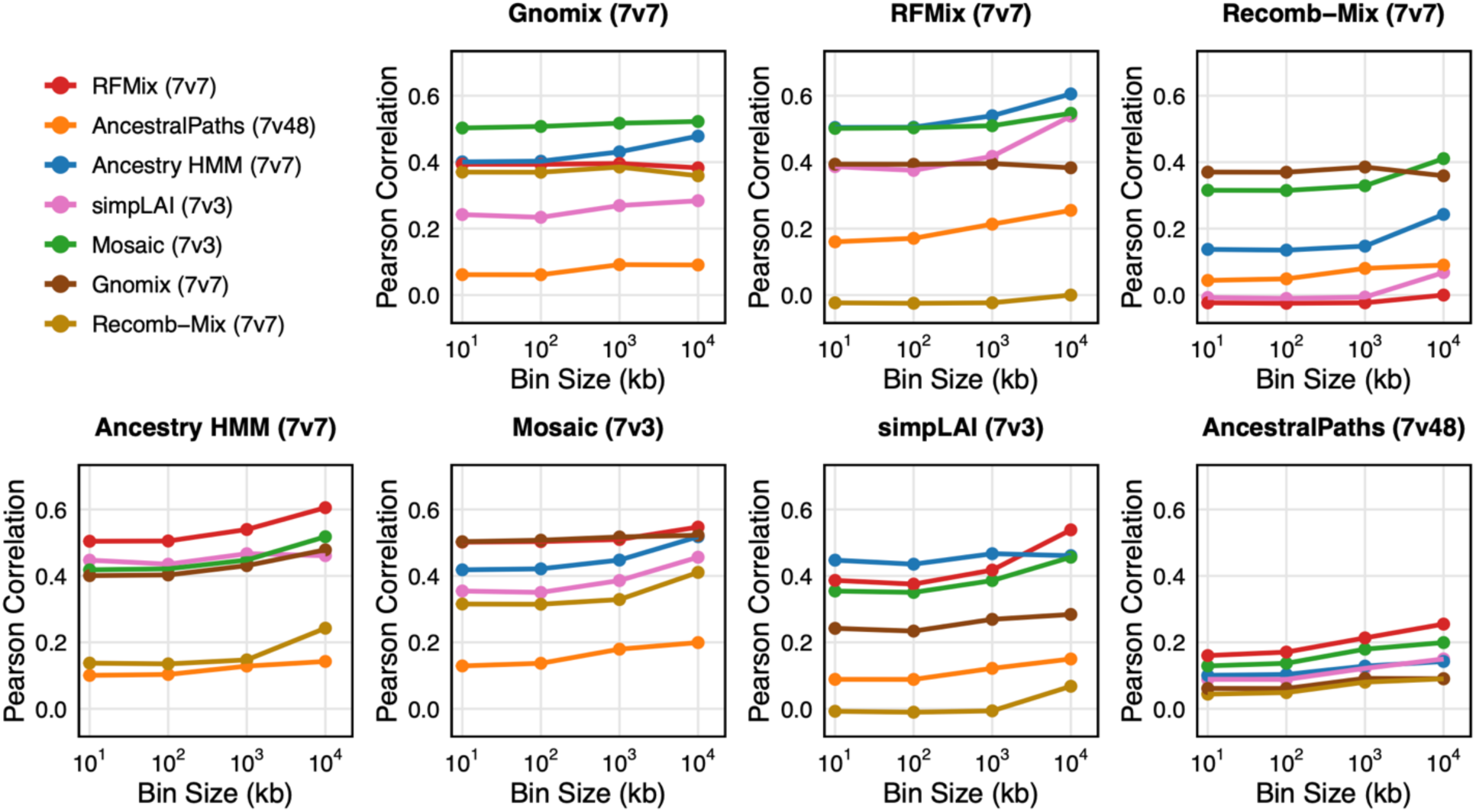
Local ancestry correlations. Pearson correlations of local ancestry proportions between LAI methods, evaluated across varying bin sizes. Source sample sizes shown in parentheses with the method title, indicating 7 farmers versus N hunter-gatherers (7vN).

In contrast, LAI methods applied to admixed African Americans from the 1000 Genomes Project showed substantially higher concordance for several approaches, with RFMix, Mosiac, and Gnomix producing pairwise genome-wide correlations > 0.9, whereas Ancestry HMM, simpLAI, and Recomb-Mix showed lower correlations (R < 0.31; Figure S8). The reduced correlations observed in the Neolithic individuals likely reflects the lower divergence between hunter-gatherer and farmer source populations, or lower data quality of the ancient DNA data.

Tract length distributions reflect inferred time since admixture, as recombination progressively breaks ancestry tracts over generations. LAI methods produced markedly different tract length distributions, with Ancestry HMM, AncestralPaths, and simpLAI produceing shorter average tracts (i.e., more ancestry switches) compared to other methods (Figures 4 and S9). Although the true tract length distribution is unknown, the maximum admixture time in these individuals is approximately 3,000 years (∼100 generations), and previous linkage disequilibrium-based estimates place average admixture events 10-30 generations before sampled admixed Neolithic populations (Lipson et al. 2017). Thus, Ancestry HMM and AncestralPaths produce implausibly short tracts, implying admixture occurring more than 100 generations before the sampled Neolithic population (Figures 4 and S9).

**Figure 4:**
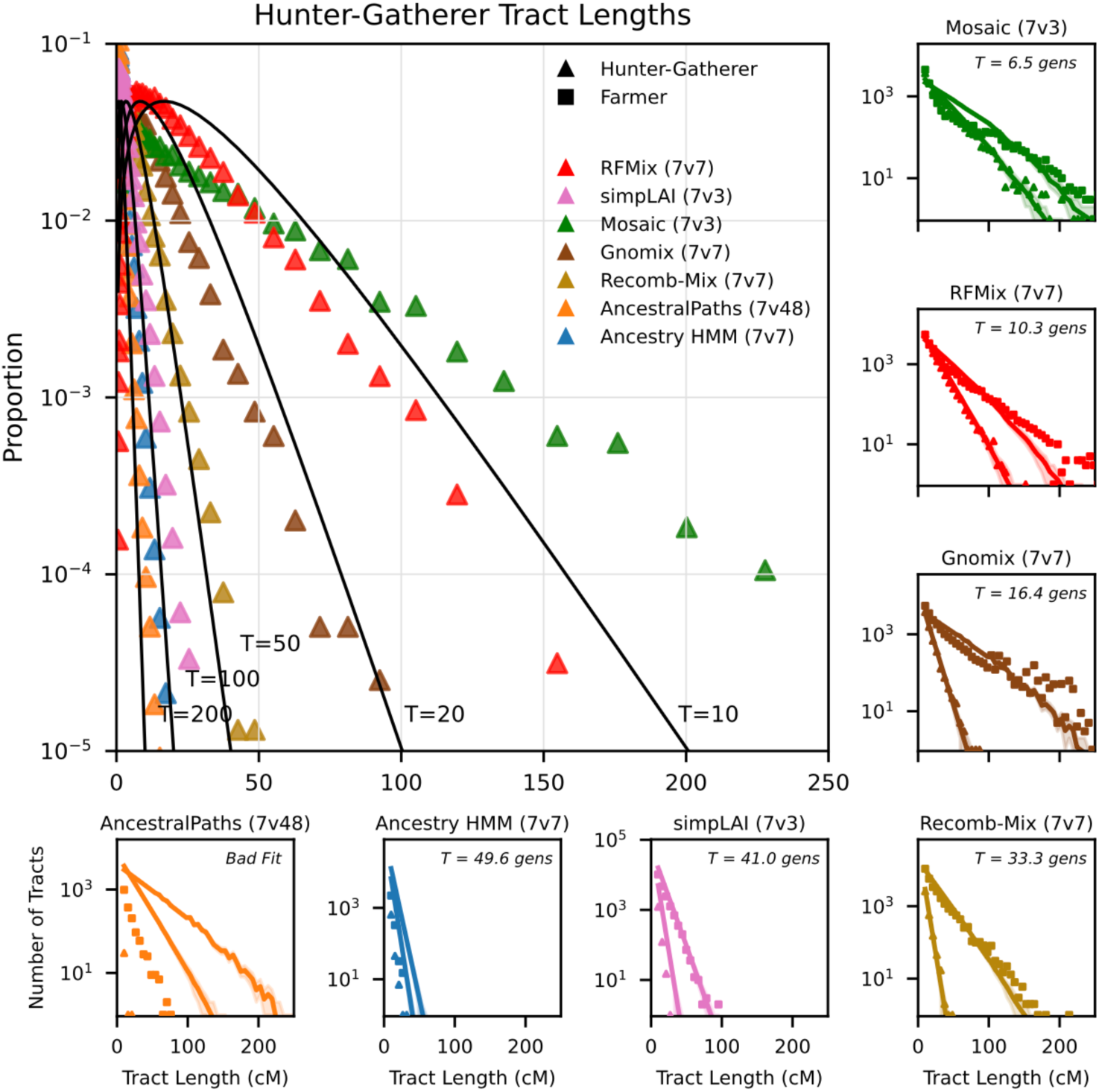
Tract length distributions by method. Proportion of hunter-gatherer ancestry tracts by length (in cM) for each method compared to theoretical single-pulse admixture models with admixture times (T) of 200, 100, 50, 20, and 10 generations and best-fit tract length distributions and inferred generations since admixture using TRACTS. Source sample sizes shown in parentheses with the method title, indicating 7 farmers versus N hunter-gatherers (7vN).

Using TRACTS (Gravel 2012), the best-fit admixture timing estimates were 10.3, 16.4, and 33.3 generations for RFMix, Gnomix, and Recomb-Mix, respectively (Figure 4). simpLAI inferred an older admixture time (41.0 generations), driven by an excess of shorter ancestry tracts in both ancestries. Mosaic inferred a more recent admixture time (6.5 generations), with tract lengths of almost full chromosomes for many individuals and very similar tract length distributions between hunter-gatherer and farmer ancestry. X chromosome tract length distributions closely matched autosomal distributions for all methods except Mosaic, which fit an admixture time of approximately 50 generations (Figure S10). Overall, RFMix and Gnomix produced tract length distributions most consistent with previous LD-based estimates of Neolithic admixture timing, with Mosaic producing relatively long tracts likely underestimating admixture timing. The remaining methods inferred shorter tracts with implausibly ancient admixture.

Consistent with the local ancestry correlation analyses, RFMix, Gnomix, and Mosaic produced highly concordant tract length distributions in African American individuals from the 1000 Genomes Project (Figure S11). In contrast, simpLAI and Ancestry HMM inferred shorter ancestry tracts, indicating bias that is due to the method rather than the data.

### Meta-analysis identifies three regions with significant deviation in local ancestry

Although methods are highly correlated at the level of global ancestry proportions, concordance across methods at the local ancestry level is modest, suggesting that each method captures partially independent signals. To identify robust deviations in local ancestry, we therefore performed a meta-analysis of Z-scores across LAI methods, accounting for correlations between methods. This approach assumes that true signals of selection should replicate across methods despite method-specific biases (Figure 5).

**Figure 5:**
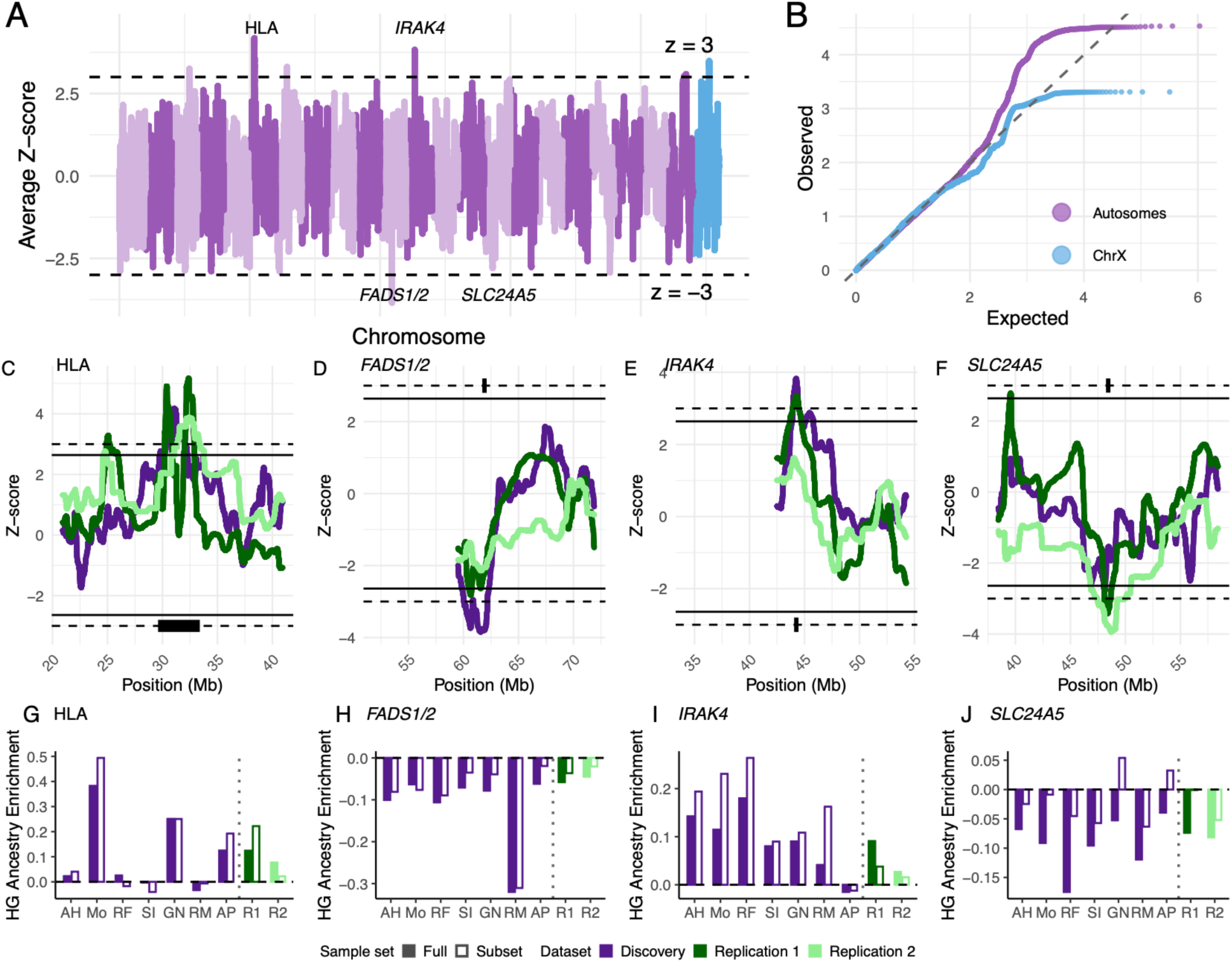
Average Z-scores across methods and replication for top hits. **A)** Meta-analyzed Z-scores for hunter-gatherer ancestry across the genome. Positive values indicate excess hunter-gatherer ancestry and negative values indicate excess farmer ancestry. Dashed lines indicate Z = 3 and Z = −3. **B)** QQ plot of Z-scores for autosomes (purple) and X chromosome (blue). **C-F)** Combined discovery and replication Z-scores for the three replicated regions and SLC24A5. Dashed lines indicate the discovery threshold (|Z| = 3), and solid lines indicate the replication threshold (|Z| = 2.64). Putative gene candidates are highlighted with black boxes. **G-J**) Absolute deviations in hunter-gatherer ancestry for each of the loci in **C-F**. Each solid bar represents the deviation for each method, and each open bar represents the deviation in a subset of (N = 27 in discovery dataset) individuals with less than 10% hunter-gatherer ancestry – we expect these individuals to be relatively unaffected by selection. Abbreviations: AH Ancestry HMM, Mo Mosaic, RF RFMix, SI simpLAI, GN Gnomix, RM Recomb-Mix, AP AncestralPaths, R1 Replication dataset 1 (Ancestry HMM), R2 Replication dataset 2 (RFMix).

Across the autosomes, six regions exceeded a threshold of |Z| > 3 (Table S1). To validate these signals, we performed LAI in two independent replication datasets: (1) 378 pseudohaploid admixed Neolithic individuals analyzed with Ancestry HMM, and (2) 1,121 imputed individuals analyzed with RFMix (Table S2). Of the six regions exceeding the discovery threshold, three replicated within 2 Mb of the lead SNP at a significance threshold of P < 0.0083 in replication dataset 1, and one of these also replicated in replication dataset 2 (Table 2). The region replicated in both replication datasets corresponds to the HLA region on chromosome 6, a previously reported ancestry outlier in admixed Neolithic populations consistent with immune-related selection (Davy et al. 2023).

**Table 2:** Significant deviations in local ancestry. Autosomal regions with significant deviations in local ancestry (|Z| > 3), showing discovery Z-scores and P-values, along with replication statistics for regions within 2 Mb of the lead SNP.

| CHR | Region | Gene | Discovery Z | Discovery P | Replication 1 Z | Replication 1 P | Replication 2 Z | Replication 2 P |
| --- | --- | --- | --- | --- | --- | --- | --- | --- |
| 6 | 31,082,285 | *HLA* | 4.17 | $2.9 \times 10^{-5}$ | 5.17 | $2.4 \times 10^{-7}$ | 3.88 | $1.0 \times 10^{-4}$ |
| 11 | 61,906,864 | *FADS1/2* | -3.66 | $2.0 \times 10^{-4}$ | -2.90 | $3.0 \times 10^{-3}$ | -2.14 | $3.2 \times 10^{-2}$ |
| 12 | 44,252,015 | *IRAK4* | 4.21 | $2.6 \times 10^{-5}$ | 3.33 | $8.6 \times 10^{-4}$ | 1.62 | $1.0 \times 10^{-1}$ |
| 15 | 48,105,416 | *SLC24A5* | -2.79 | $5.0 \times 10^{-3}$ | -3.60 | $3.0 \times 10^{-4}$ | -3.94 | $8.1 \times 10^{-5}$ |

The second region replicated in replication dataset 1 occurs at the *FADS1/2* locus on chromosome 11, which has been implicated in dietary adaptation in ancient European populations (Mathieson and Mathieson 2018; Irving-Pease et al. 2024). The third replicated region on chromosome 12 also overlaps multiple genes, with *IRAK4*, involved in innate immunity, representing a possible candidate (Suzuki et al. 2006). One region exceeds |Z| > 3 on the X chromosome discovery dataset but did not replicate at P < 0.05 in replication dataset 1, the only replication dataset with X chromosome data (Table S3; Figures S12 and S13).

Another region previously identified as an ancestry outlier in admixed Neolithic populations, *SLC24A5* on chromosome 15, was not identified as an ancestry outlier in the discovery dataset but passes |Z| > 3 in both replication datasets (Figure 5F; Davy et al. 2023).

The replicated Z-score deviations are directionally consistent across methods and robust to variation in source sample size (Tables S4-S6; Figures S14–S17). To further validate these signals, we compared ancestry deviations across methods and replication datasets, analyzing both the full sample and a subset of 27 individuals with <10% hunter-gatherer ancestry. These older individuals (mean age: 7,127 years before present) are closer to the initial admixture and thus expected to exhibit smaller ancestry deviations due to limited time for selection to act.

We find ancestry deviations at *SLC24A5* and *FADS1/2* are consistent across methods and lower in the subset of low-ancestry individuals – consistent with reflecting the effect of selection. Deviation at *IRAK4* is consistent across methods but high in the low-ancestry individuals, suggesting that it might reflect ancestry misclassification rather than the effects of selection. Finally, deviation at the HLA is inconsistent across methods and replication datasets and high in the low-ancestry individuals. In summary, ancestry deviations at *SLC24A5* and *FADS1/2* likely indicate selection at those loci while other deviations, particularly at the HLA, likely indicate that ancestry is being systematically misclassified at those loci.

## Discussion

Our results demonstrate both the utility and limitations of applying local ancestry inference to ancient genomes. By systematically evaluating multiple LAI approaches, we show that existing methods are highly sensitive to source composition and technical aspects of data preparation.

These are likely exacerbated by features of ancient DNA, including limited reference panels, temporal divergence between sources and admixed individuals, and genotype uncertainty.

Despite these biases, integrating results across methods recovers known adaptive loci and identifies additional candidates. Together, these findings show that method-aware, multi-method approaches can yield robust evolutionary insights, while highlighting that unvalidated LAI-based scans may misestimate the magnitude or even direction of ancestry-specific selection.

The replication of Neolithic selection signal at *FADS1* provides biological validation of this framework. *FADS1*, involved in fatty acid metabolism, has been repeatedly implicated in dietary adaptation in Neolithic and Bronze Age Europeans (Mathieson and Mathieson 2018; Irving-Pease et al. 2024). Although *SLC24A5* was not identified in the discovery meta-analysis, it is a well-characterized pigmentation gene, with variants associated with lighter skin pigmentation having undergone strong selection in European populations over the past several thousand years (Lamason et al. 2005; Grossman et al. 2010; Ju and Mathieson 2021). Both *FADS1* and *SLC24A5* show consistent deviations in Neolithic populations across methods and datasets. Importantly, local ancestry deviation reflects not only post-admixture selection but also pre-existing differentiation between source populations. Thus, loci such as *FADS1* and *SLC24A5* likely reflect adaptive differences associated with agricultural lifestyles. The consistently lower ancestry deviation observed in older individuals across methods and datasets confirms the robustness of these signals to methodological and data-type variation.

Indeed, frequency-based analyses also identify *FADS1* and *SLC24A5* as selection signals. Based on known causal SNPs (*FADS1*, rs174546; *SLC24A5*, rs1426654), we see observed change in frequency of 0.23 for *FADS1* and 0.33 for *SLC24A5* (Akbari et al. 2026). However, if the expected allele frequency change is calculated from the difference in hunter-gatherer and farmer allele frequencies and the observed change in ancestry proportions, the expected change is only 0.03 for *FADS1* and 0.06 for *SLC24A5*. This suggests that most of the allele frequency change is due to change within the same ancestry background rather than shifts in local ancestry, or that the shifts in local ancestry are systematically underestimated.

Beyond these established loci, we identify *IRAK4* as a novel candidate region. *IRAK4* encodes a key kinase in innate immune signaling pathways critical for host defense against pathogens (Suzuki et al. 2006). Its signal aligns with immune-related adaptation, potentially reflecting selection on hunter-gatherer–introduced variation responding to local pathogen environments.

However, ancestry at *IRAK4* shows substantial deviation even in individuals with low (<10%) hunter-gatherer ancestry, suggesting that these are the result of ancestry misclassification. This misclassification might represent a technical artefact or a more complicated form of selection. For example, if our source individuals are mismatched to the true source populations, and selection occurred between the true sources and our sample.

We also replicated a previously reported excess of hunter-gatherer ancestry at the HLA (Davy et al. 2023; da Silva et al. 2025). However, this signal was not consistent across methods and showed evidence of ancestry misclassification. On one hand, the HLA plays a central role in immune function and is expected to be a frequent target of selection. Indeed, it is often identified as a target of both balancing and directional selection, including in admixed populations (Prugnolle et al. 2005; Meyer et al. 2018; da Silva et al. 2025). On the other hand, LAI at this locus is particularly challenging and signals of directional selection in present-day admixed populations seem to be largely artefactual (Bhatia et al. 2014; Mandla et al., forthcoming). We conclude that local ancestry at the HLA may be too unreliable to make reliable inference about natural selection or trait associations (Davy et al. 2023; Barrie et al. 2024).

The X chromosome is often excluded from LAI, and selection analyses more broadly, yet may harbor extensive and important signals of selection (Skov et al. 2023). We identify one region exceeding the |Z| > 3 threshold on the X chromosome (X:97.9 Mb), although it does not replicate in an independent dataset (Figure S12). The lack of replication may reflect limited power, as the pseudohaploid X chromosome dataset contains only 2,061 SNPs. The X:97.9 Mb region shows excess hunter-gatherer ancestry, and nearby genes include *LINC03077* and *DIAPH2*. While no genes directly within this region have been previously identified as selection targets, nearby loci have been implicated in selection scans, suggesting this region warrants further investigation with higher-resolution data (Villegas-Mirón et al. 2021; Skov et al. 2023).

A limitation of this study is the absence of ground truth local ancestry in admixed Neolithic genomes. Consequently, our benchmarking evaluates concordance between methods and agreement with independent expectations, including genome-wide ancestry estimates, tract length distributions, admixture time estimates, and replication of selection signals, rather than local ancestry accuracy. Simulation studies can provide a complementary framework by enabling direct assessment against known ancestry tracts. Indeed, while our study was under review, a benchmarking study using simulations designed to reflect ancient DNA conditions reported broadly similar conclusions, including superior performance of RFMix across several evaluation metrics (Bougiouri et al., unpublished data). Further, they report that FLARE frequently infers 100% major ancestry when run with small source sample sizes. In contrast to our results, they find Mosaic underestimates and simpLAI overestimates tract lengths. These results suggest that empirical and simulation-based benchmarking provide complementary evidence for method performance, with empirical analyses capturing complexities of real ancient genomes that remain difficult to simulate.

Our results translate into guidelines for future studies applying LAI to ancient DNA. We recommend validating findings across multiple methods and benchmarking against global ancestry estimates (e.g., qpAdm and ADMIXTURE). Evaluating concordance in both global and local ancestry, as well as comparing tract length distributions to theoretical expectations, can help to identify method-specific biases. Among the methods evaluated here, we recommend Gnomix and RFMix, and possibly Mosaic. These methods produced the most accurate ancestry proportions and plausible tract length distributions. All other methods produced biased estimates and underestimated tract lengths. An important factor may be method-internal rephasing (RFMix, Gnomix, Mosiac, Recomb-Mix; Table 1) which corrects phasing errors and therefore increases length of tracts, while post-LAI rephasing appears to make little difference.

Additionally, we use a simplified single pulse admixture model which we believe fits a weighted average time since admixture for all individuals but given the evidence for multi-pulse admixture in the Neolithic, future work should expand this analysis to admixture under multi-pulse models (Lipson et al. 2017; Irving-Pease et al. 2024).

These findings are particularly relevant for studies of ancient populations with limited sample sizes. Although larger ancient DNA datasets are rapidly becoming available, many populations remain sparsely sampled. In such settings, biases in LAI may be amplified due to small and unbalanced reference panels and reduced statistical power. Systematic benchmarking, as performed here, is therefore essential for ensuring robust inference. More broadly, our results demonstrate the importance of method validation when extending tools developed for present-day populations to ancient genomic data and provide an empirical framework for studying evaluating LAI in challenging datasets.

## Methods

### Data and data processing

We analyzed autosomal and X chromosome ancient DNA data from the imputed and pseudohaploid discovery dataset (Gamba et al. 2014; Lazaridis et al. 2014; Olalde et al. 2014; Skoglund et al. 2014; Allentoft et al. 2015; Günther et al. 2015; Olalde et al. 2015; Cassidy et al. 2016; Kılınç et al. 2016; González-Fortes et al. 2017; Martiniano et al. 2017; Fernandes et al. 2018; Valdiosera et al. 2018; Antonio et al. 2019; Brace et al. 2019; González-Fortes et al. 2019; Malmström et al. 2019; Sánchez-Quinto et al. 2019; Schroeder et al. 2019; Brunel et al. 2020; Scorrano et al. 2022; Allentoft et al. 2024). All analyses were performed using the GRCh37 (hg19) reference genome. Discovery dataset autosomal variant call format (VCF) files for shotgun-sequenced ancient individuals are publicly available at https://erda.ku.dk/archives/917f1ac64148c3800ab7baa29402d088/published-archive.html. Imputation was previously performed using GLIMPSE (Rubinacci et al. 2021) with the 1000 Genomes Project (Auton et al. 2015) phase 3 reference panel (see methods described in Allentoft et al., 2024).

The discovery dataset was taken from publicly available VCF files reported in Allentoft et al. (2024), filtered for coverage >0.1, relatedness, and genome-wide average imputation genotype probability less than 0.98 (see Allentoft et al. (2024) Methods and Supplementary Notes). We selected individuals based on IBD cluster groupings from Supplementary Data VII, selecting admixed Neolithic farmers (Farmer_Europe_early, Farmer_EuropeW_late, Farmer_EuropeE_late), Western hunter-gatherers (HG_EuropeW), and Anatolian farmers (WesternAsia subset of Farmer_Europe_early). SNPs were filtered to retain only those overlapping between imputed VCF data and pseudohaploid array SNPs, with minor allele frequency (MAF) > 0.01 and imputation quality (INFO score) > 0.8, resulting in 572,223 autosomal SNPs.

For the X chromosome discovery dataset, we downloaded published sequence data from the same 176 admixed Neolithic, 7 farmer, and 48 hunter-gatherer individuals listed in Allentoft et al. (2024) Supplementary Data VII. Where BAM files were unavailable, we aligned raw FASTQ files or merged multiple libraries. We used SAMtools (Li et al. 2009) to extract X chromosome reads and applied GLIMPSE (Rubinacci et al. 2021) with the 1000 Genomes Project (Auton et al. 2015) reference panel for imputation. Only the non-pseudoautosomal regions (non-PAR) of the X chromosome (GRCh37/hg19 coordinates X: 2,699,520–154,931,044) were retained. We filtered X chromosome SNPs for MAF > 0.05 across admixed individuals, source populations, and the 1000 Genomes Project (Auton et al. 2015) reference panel, resulting in 157,610 SNPs.

Replication dataset 1 consisted of an independent dataset of 378 admixed Neolithic individuals, 98 hunter-gatherers, and 48 farmers extracted from the Allen Ancient DNA Resource (AADR) 1240k dataset (Skoglund et al. 2012; Mathieson et al. 2015; Haak et al. 2015; Lazaridis et al. 2016; Broushaki et al. 2016; Hofmanová et al. 2016; Omrak et al. 2016; Jones et al. 2017; Kılınç et al. 2017; Günther et al. 2018; Olalde et al. 2018; Mathieson et al. 2018; Villalba-Mouco et al. 2019; Furtwängler et al. 2020; Rivollat et al. 2020; Immel et al. 2021; Yaka et al. 2021; Marchi et al. 2022; Mallick et al. 2024). This dataset is the dataset analyzed by Davy et al. (2023) with the individuals in our discovery dataset removed. The datasets were merged in PLINK, and pairwise identity-by-descent (IBD) was calculated; all individuals in pairs with PI_HAT > 0.9 were removed to exclude duplicate or near-identical individuals across datasets. We additionally removed individuals with identical names between datasets. We filtered SNPs as the intersection with the discovery SNP sets for the autosomes, yielding 572,223 autosomal SNPs and included all 2,061 reported X chromosome SNPs.

Replication dataset 2 (Lipson et al. 2017; Fregel et al. 2018; Feldman et al. 2019; Narasimhan et al. 2019; Olalde et al. 2019; Scheib et al. 2019; Cassidy et al. 2020; Fernandes et al. 2020; Gokhman et al. 2020; Marcus et al. 2020; Freilich et al. 2021; Harney et al. 2021; Novak et al. 2021; Papac et al. 2021; Saupe et al. 2021; Seguin-Orlando et al. 2021; Villalba-Mouco et al. 2021; Altınışık et al. 2022; Ariano et al. 2022; Arzelier et al. 2022; Childebayeva et al. 2022; Dulias et al. 2022; Fowler et al. 2022; Lazaridis et al. 2022; Maróti et al. 2022; Patterson et al. 2022; Rohland et al. 2022; Yu et al. 2022; Chyleński et al. 2023; Koptekin et al. 2023; Mattila et al. 2023; Penske et al. 2023; Posth et al. 2023; Rivollat et al. 2023; Skourtanioti et al. 2023; Simões et al. 2023; Wang et al. 2023; Arzelier et al. 2024; Gelabert et al. 2024; Seersholm et al. 2024; Simões et al. 2024; Lazaridis et al. 2025; Szécsényi-Nagy et al. 2025; Akbari et al. 2026; Olalde et al. 2026) included 1,121 admixed Neolithic individuals, 32 farmers, and 26 hunter-gatherers reported by Akbari et al. (2026). Data are publicly available from https://dataverse.harvard.edu/dataset.xhtml?persistentId=doi:10.7910/DVN/7RVV9N. The dataset is comprised of a mixture of shotgun and capture sequencing data and had been imputed using GLIMPSE. Individuals were selected based on geographic distribution (hunter-gatherers: northern, southwestern, central, and southeastern Europe; farmers: southeastern Europe; admixed Neolithic: northern, southwestern, central, and southeastern Europe), published qpAdm ancestry estimates (hunter-gatherers: WHG > 0.9; farmers: ANF > 0.95; admixed Neolithic: 0 < WHG < 0.7, ANF > 0.3, and (ICR + EHG) < 0.1), and sample age (hunter-gatherers: 5,000–12,000 years before present; farmers: 7,000–11,000; admixed Neolithic: 3,000–10,000). qpAdm estimates were obtained from the accompanying ‘Ancestry composition estimates (qpAdm models)’ (Ancestry_Composition_4way_qpAdm_Fernandes2020.tsv). We also removed any individuals that were included in either the discovery dataset, or replication dataset 1. SNPs were then filtered to the intersection with the same 572,223 autosomal SNP set used in the discovery analysis.

### Local ancestry inference on autosomes

We ran six LAI methods (Ancestry HMM, Mosaic, RFMix, simpLAI, Gnomix, Recomb-Mix) on the extracted 176 admixed individuals, SNP sets, and source populations, and extracted published calls for AncestralPaths from published VCF files (https://erda.ku.dk/archives/917f1ac64148c3800ab7baa29402d088/publishedarchive.html; Allentoft et al. 2024).

AncestralPaths published local ancestry calls were extracted for the same 176 admixed individuals present in our discovery dataset. Unlike the other LAI methods benchmarked here, AncestralPaths does not use a fixed reference panel: its classifier is trained on simulated tree sequences under a demographic model, then applied to a RELATE tree sequence constructed from real samples. The 7 farmer and 48 hunter-gatherer sample counts reported for our discovery dataset (Allentoft et al. 2024, Supplementary Data VII) describe the real sample composition available for this comparison, not a confirmed reference panel size used internally by AncestralPaths.

AncestralPaths employs a neural network trained on simulated European demographic histories to assign a discrete path value (1–6) at each SNP, corresponding to population labels. For admixed Neolithic populations, calls included ancestries beyond hunter-gatherers and farmers; we recoded these to their closest source population based on phylogenetic proximity in the training data (WHG: 3; ANA: 1; 2, 5, 6 → 1; 4 → 3). We extracted the 572,223 autosomal SNPs from AncestralPaths VCF calls. To address the observed inflated hunter-gatherer ancestry estimates on chromosomes 6 and 9, we adjusted SNP-level means per chromosome by the difference between the chromosome mean and the genome-wide mean at the population level (Figure S1).

Ancestry HMM allele frequencies were calculated with plink/1.90 Beta 6.18 for hunter-gatherer, farmer, and admixed individuals. Input data were separated by haplotype for each individual to preserve linkage disequilibrium (LD) information. We used parameters of 35 generations since admixture, effective population size (Ne) of 10,000, and admixture proportions of 0.8 farmers and 0.2 hunter-gatherers.

We performed Mosaic analyses using bcftools/1.20 to prepare genotype files and ran with identical admixture parameters as Ancestry HMM (35 generations, Ne=10,000).

We implemented RFMix with the PopPhased option, assuming 35 generations since admixture and setting -n 5 for the minimum number of reference haplotypes per tree node to accommodate unbalanced reference sample sizes. simpLAI was run using parameters recommended in its documentation (https://github.com/CMPG/simpLAI/blob/main/README.md): -s 1e6, -I 5e5, -n 2000, -m 1000, -t 5.

We ran Gnomix by chromosome using default parameters from its documentation (https://github.com/AI-sandbox/gnomix/blob/main/README.md) including the default config.yaml file.

Similarly, we ran Recomb-Mix by chromosome using included GRCh37 genetic maps, final allele frequency filter of > 1% (-f 0.01) and all other default parameters (https://github.com/ucfcbb/Recomb-Mix/blob/main/README.md).

For uniform comparison, we converted all methods’ SNP-level calls to RFMix format. For methods producing ancestry calls in SNP windows (Mosaic, simpLAI, Gnomix, Recomb-Mix), we extended each window’s call to all SNPs contained within. To correct for phasing errors, we applied TRACTOR rephasing (unkink_2way_mspfile.py from https://github.com/Atkinson-Lab/Tractor) to each LAI output, including those which perform internal rephasing (Table 1).

To evaluate the impact of reference panel size, we down sampled hunter-gatherer source individual sample sizes from 48 to 7, 3, and 1 for all methods except AncestralPaths. For RFMix, Gnomix, Ancestry HMM, and Mosaic, we filtered the data to include only inferred local ancestry calls with posterior probability > 0.9 at each down sampled level.

For LAI methods, source sample sizes consisted of combinations of 7 farmers (full farmer sample size) and hunter-gatherer sample sizes of: 48, 7, 3, and 1 to test for biases in source sample size imbalance. AncestralPaths published calls were produced with source sample sizes of 7 farmers and 48 hunter-gatherers. Sources were randomly sampled to obtain smaller sample sizes (Table S7). We selected the optimal source sample sizes to use for each method by optimizing for best fit with qpAdm using Pearson’s correlation between individuals’ global ancestry with qpAdm, slope of regression line with qpAdm, and mean difference with qpAdm individual estimates (Figures 2 and S2). For methods that reported posterior probabilities for ancestry calls, filtering on > 0.9 posterior probability further improved fit of LAI results to qpAdm and thus were selected as optimal results (Figure S2).

### Benchmarking local ancestry inference methods

We estimated genome-wide hunter-gatherer ancestry proportions per admixed individual by dividing the number of SNPs assigned hunter-gatherer ancestry by the total number of SNPs (572,223). We assessed correlations among ancestry estimates from LAI methods, qpAdm (Haak et al. 2015), and ADMIXTURE (Alexander et al. 2009) with Pearson and Spearman’s correlation coefficients across individuals.

We ran qpAdm and unsupervised ADMIXTURE (K = 2) with consistent reference and admixed individuals, and SNP set. qpAdm analyses were performed on both pooled admixed individuals and individually using outgroup populations from the AADR. Outgroup populations included “Karitiana.DG”, “Mbuti.DG”, “Papuan.DG”, “Han.DG”, “Russia_Ust_Ishim_HG_published.DG”, “Belgium_UP_GoyetQ116_1_published_all”, “Russia_MA1_HG.SG”, “Russia_Kostenki14.SG”, “Italy_North_Villabruna_HG”, “Czech_Vestonice16”, “Russia_Kostenki14”, “Spain_ElMiron”, “Russia_AfontovaGora3”. qpAdm and ADMIXTURE were run on both imputed and pseudohaploid data (Figure S4).

To assess sample size sensitivity, we ran unsupervised ADMIXTURE (K = 2) with different hunter-gatherer sample sizes (7v48, 7, 3, and 1). Regression analyses between qpAdm ancestry estimates and each LAI method’s estimates were performed to calculate slope coefficients and mean differences. Optimal source sample sizes were selected by minimizing average ancestry estimate deviation from qpAdm and maximizing correlation coefficient. We further calculated mean squared error between ancestry estimates and qpAdm to order methods in Figures 2-4.

Local ancestry correlations were computed between all pairs of LAI methods at the population level by taking the average hunter-gatherer ancestry proportion for all 176 individuals in genomic bins of 10 kb, 100 kb, 1 Mb, and 100 Mb and computing the Pearson correlation. Similarly, at the individual level, we computed Pearson correlation of inferred hunter-gatherer ancestry proportion for each individual for each pair of methods in 100 kb bins and averaged the correlation across pairs of methods. (Figure S7).

### Sensitivity analyses (source size, imputation, phasing)

We tested LAI methods (Ancestry HMM, simpLAI, Mosaic, RFMix, Gnomix, Recomb-Mix) on 1000 Genomes Project(Auton et al. 2015) data with admixed population African American (ASW, n=61), and source populations Yoruba (YRI, n=7) and European (CEU, n=48) populations. We ran unsupervised ADMIXTURE (K = 2) and qpAdm using an expanded set of outgroups (GBR, FIN, CHS, CDX, IBS, KHV, PJL, GWD, ESN, BEB, MSL, STU, ITU, JPT, LWK, GIH). Following the admixed Neolithic analyses, we compared global ancestry estimates across methods and local ancestry correlations to test if the source sample sizes were biasing LAI results in favor of the minor ancestry or ancestry with a larger source sample size (Figure S3).

Similarly, we analyzed tract length distributions for each method following from *Tract length analysis* Methods section.

We assessed biases introduced by imputation by running qpAdm, ADMIXTURE, and Ancestry HMM on unimputed Allentoft VCFs and compared global ancestry estimates to those from imputed data, using matched SNP counts and sources (7v48; Figure S4).

We addressed issues with phasing by running LAI methods on the X chromosome, where we ran LAI on X for males and XX for females (see *X chromosome* methods section below). We computed ancestry estimates on the X with qpAdm, ADMIXTURE, and LAI methods RFMix, simpLAI, Mosaic, Gnomix, Recomb-Mix and Ancestry HMM and compared ancestry proportion of males vs females to examine biases introduced by phasing on females by running a two-sided T-test for difference between male and female average hunter gatherer ancestry on the X chromosome (Figure S5). Similarly, we compared tract length distributions between males and females by computing tract lengths of each and then analyzing using QQ analysis of log10 tract lengths (cM) for both farmer and hunter-gatherer ancestry (Figure S6; following from *Tract length analysis* Methods section below).

### Tract length analysis

For each individual and chromosome, we computed the lengths of continuous ancestry tracts by counting the cM distance of consecutive SNPs assigned to the same ancestry before switching. Separate tract length distributions were generated for hunter-gatherer and farmer ancestries.

We compared empirical tract length distributions to theoretical expectations using the formula:

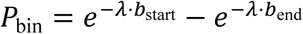

where 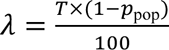, with *T* as the number of generations since admixture and *p*pop as the admixture proportion.

Using TRACTS(Gravel 2012), we inferred admixture timing assuming a single pulse model with input parameters of 35 generations, hunter-gatherer ancestry proportion *R* and time scaling factor of 100. Tracts shorter than 10 centimorgans (cM) were excluded to reduce noise from phasing errors and genotype uncertainty. Visualization was performed using the fancyplotting.py script from the TRACTS package.

### X chromosome analysis

We applied unsupervised ADMIXTURE (K = 2), qpAdm, and LAI methods (RFMix, simpLAI, Mosaic, Ancestry HMM, Gnomix, Recomb-Mix) to X chromosome data with consistent source sample sizes determined for autosomes (7 hunter-gatherers versus N farmers).

For Ancestry HMM, we encoded females as two haplotypes and designated as ploidy 2, and males as one haplotype with ploidy set to 1. For methods designed for diploid data (Mosaic, RFMix, simpLAI, Gnomix, Recomb-Mix), we treated males as pseudo-diploid by duplicating their haploid sequence(Browning et al. 2016). Post LAI, one haplotype per male was removed to avoid double counting in following analyses.

We compared global ancestry proportions between males and females using two-sample t-tests. Differences in ancestry tract length distributions by sex were assessed via quantile-quantile analysis of log10-transformed tract lengths (cM) stratified by ancestry.

### Ancestry deviation analysis

We computed standardized Z-scores for each LAI method at the population level by subtracting the genome-wide mean hunter-gatherer ancestry from each SNP’s mean ancestry and dividing by the genome-wide standard deviation of ancestry within sliding windows of 51 SNPs (step size = 1 SNP).

To integrate evidence across methods, Z-scores were combined within sliding windows of 51 SNPs (step size = 1 SNP). We accounted for correlation among methods by estimating the covariance matrix of method-specific Z-scores genome-wide. The combined Z-score was calculated as:

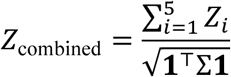

where Σ is the covariance matrix and **1** is a vector of ones.

Regions of long-range linkage disequilibrium (n=22) and 2.5 Mb flanking unmappable regions were excluded prior to Z-score calculation(Price et al. 2008). We used source sample sizes of 7 hunter-gatherers and 48 farmers for all methods except simpLAI, which had matched reference sample sizes of 7 for both sources to increase power (Figures 5 and S14).

The initial discovery significance threshold was set at ∣ *Z* ∣ > 3. We calculated autosomal and X chromosome Z-scores separately.

### Replication analyses

For replication dataset 1, we performed Ancestry HMM on pseudohaploid autosomal and X chromosome datasets consisting of 378 admixed Neolithic individuals separately. Replication Z-scores were computed in 51-SNP sliding windows with step size 1.

For replication dataset 2, we ran RFMix V1.9 with the same input parameters as the discovery dataset (with the PopPhased option, assuming 35 generations since admixture) and computed Z-scores in 51-SNP sliding windows with step size 1 after removing regions of long-range linkage disequilibrium (n=22) and 2.5 Mb flanking unmappable regions(Price et al. 2008).

For autosomal loci, a Bonferroni-corrected significance threshold of *p* = 0.05/10 hits corresponded to ∣ *Z* ∣ = 2.64 within a 2mb region flanking each discovery hit. For the X chromosome, a threshold of *p* = 0.05 corresponded to ∣ *Z* ∣= 1.6.

We compared ancestry deviations in these regions across LAI methods and for autosomes, across source sample sizes of 7vN and RFMix 7v7, to confirm direction of signals replications (Figures S14-S17, Tables S4-S6).

To evaluate ancestry deviations as a function of time since admixture, we defined subsets of individuals with low hunter-gatherer ancestry using qpAdm estimates. In the discovery dataset, we filtered 176 admixed individuals for <0.1 hunter-gatherer ancestry, resulting in 27 individuals (mean age: 7,127 years before present).

For each LAI method (RFMix, simpLAI, Mosaic, Gnomix, Recomb-Mix, Ancestry HMM, and AncestralPaths), we calculated genome-wide average ancestry across all 176 individuals and separately for the subset of 27 low hunter-gatherer ancestry individuals (Table S7). For each of the four replicated loci (HLA: chr6:30.0–33.0 Mb; FADS1/2: chr11:59.0–65.0 Mb; IRAK4: chr12:42.25–46.25 Mb; SLC24A5: chr15:46.11–50.11 Mb; hg19), ancestry deviations were calculated as the difference between local ancestry at the locus and the corresponding genome-wide average. Method results follow from meta-analysis source sample sizes (7v48), besides simpLAI (7v7).

We applied the same procedure to replication dataset 1 by selecting the 30 individuals with the lowest hunter-gatherer ancestry and to replication dataset 2 by selecting 692 admixed Neolithic individuals with qpAdm estimates <0.1 hunter-gatherer ancestry. In both cases these individuals were used to define the low ancestry subset. Ancestry deviations were compared between full datasets and low hunter-gatherer ancestry subsets across methods and datasets.

We compared the difference between expected and observed allele frequency changes for *FADS1* and *SLC24A5* using rs174546 for *FADS1* and rs1426654 for *SLC24A5* from ***AGES: A****ncient **GE**nome **S**election* (https://reich-ages.rc.hms.harvard.edu/#/). We estimated the expected allele frequency change attributable to ancestry shifts as:

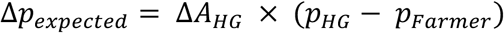

where Δ*A_HG_* is the average change in hunter-gatherer ancestry across the six LAI methods and *p_HG_* and *p_Farmer_* are the allele frequencies in the hunter-gatherer and farmer source populations reported by AGES. We compared these expected changes with the observed allele frequency changes reported by AGES between 8.6 kya and 5 kya for rs174546 (FADS1) and rs1426654 (SLC24A5).

## Code and data availability

Code to reproduce the analysis in this paper is available at https://github.com/gmies/LAI_neolithic/. Data to generate main text figures, supplemental text figures, and LAI calls are available at https://zenodo.org/records/21921239.

## Acknowledgments

This work was supported in part by the National Institute of General Medical Sciences R35GM133708 and T32GM156697. The content is solely the responsibility of the authors and does not necessarily represent the official views of the NIH. We thank Ziyue Gao for helpful discussions about the manuscript.

