## Supplemental Figures for "Local ancestry inference identifies robust evidence of selection in Neolithic Europe"

### Supplementary Figures

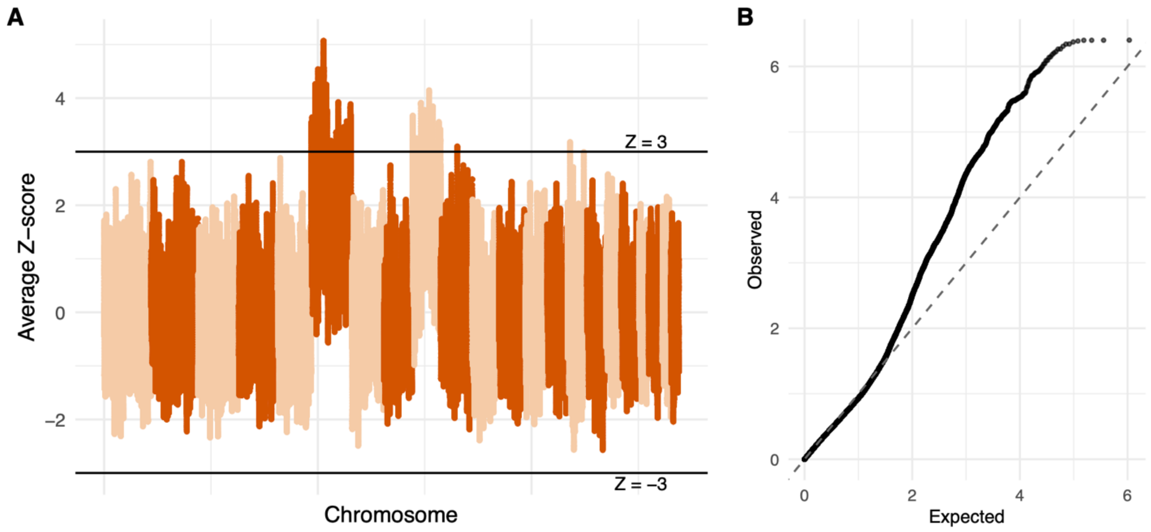


**Figure S1. AncestralPaths Z-scores for discovery dataset (N = 176). A)** Manhattan plot showing elevated hunter-gatherer ancestry on chromosomes 6 and 9 prior to per-chromosome normalization to the genome-wide average. **B)** QQ plot of Z-scores.

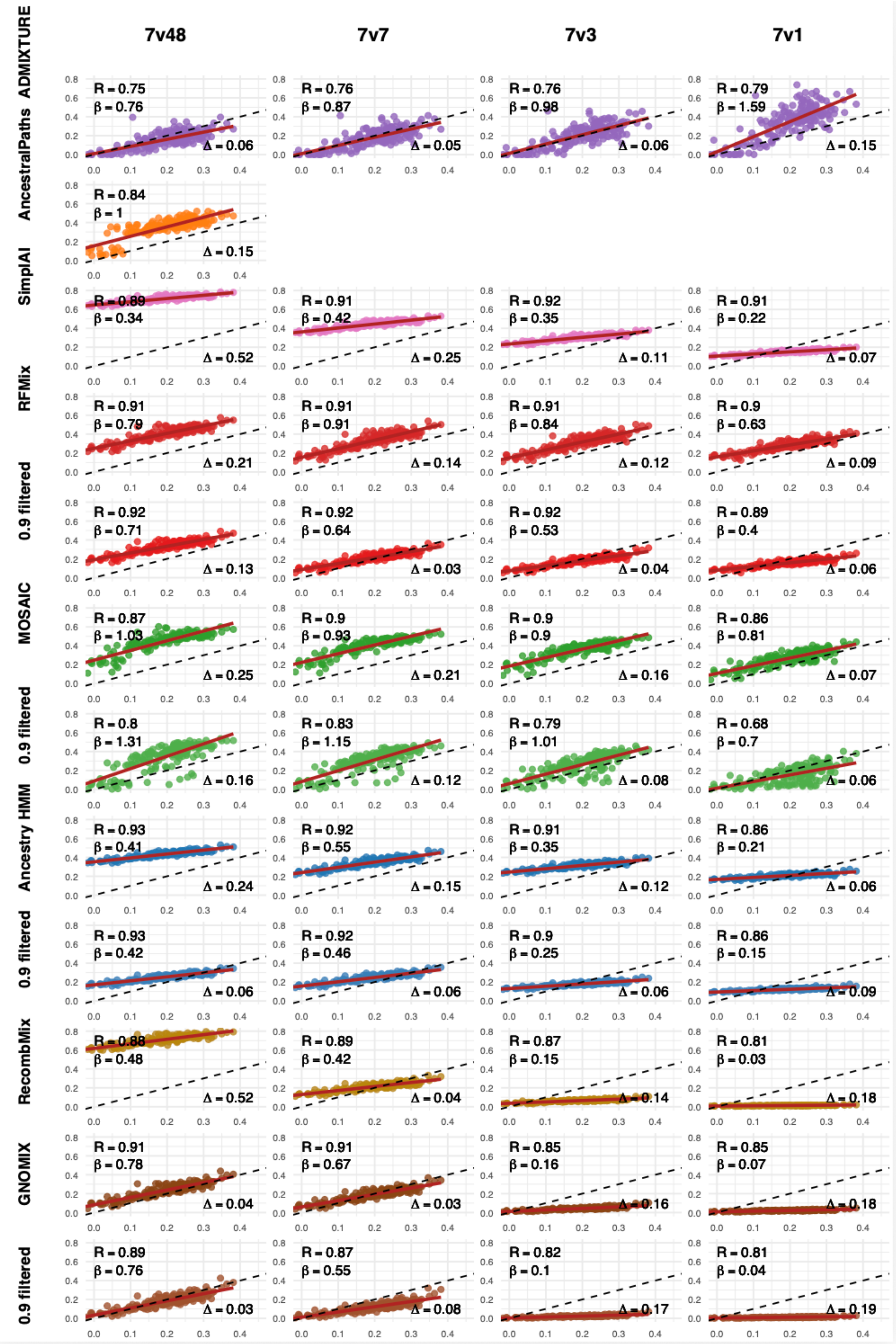

7v48
7v7
7v3
7v1
ADMIXTURE
AncestralPaths
SimplAI
RFMix
0.9 filtered
MOSAIC
0.9 filtered
Ancestry HMM
0.9 filtered
RecombMix
GNOMIX
0.9 filtered
R = 0.75 β = 0.76 Δ = 0.06
R = 0.76 β = 0.87 Δ = 0.05
R = 0.76 β = 0.98 Δ = 0.06
R = 0.79 β = 1.59 Δ = 0.15
R = 0.84 β = 1 Δ = 0.15
R = 0.89 β = 0.34 Δ = 0.52
R = 0.91 β = 0.42 Δ = 0.25
R = 0.92 β = 0.35 Δ = 0.11
R = 0.91 β = 0.22 Δ = 0.07
R = 0.91 β = 0.79 Δ = 0.21
R = 0.91 β = 0.91 Δ = 0.14
R = 0.91 β = 0.84 Δ = 0.12
R = 0.9 β = 0.63 Δ = 0.09
R = 0.92 β = 0.71 Δ = 0.13
R = 0.92 β = 0.64 Δ = 0.03
R = 0.92 β = 0.53 Δ = 0.04
R = 0.89 β = 0.4 Δ = 0.06
R = 0.87 β = 1.03 Δ = 0.25
R = 0.9 β = 0.93 Δ = 0.21
R = 0.9 β = 0.9 Δ = 0.16
R = 0.86 β = 0.81 Δ = 0.07
R = 0.8 β = 1.31 Δ = 0.16
R = 0.83 β = 1.15 Δ = 0.12
R = 0.79 β = 1.01 Δ = 0.08
R = 0.68 β = 0.7 Δ = 0.06
R = 0.93 β = 0.41 Δ = 0.24
R = 0.92 β = 0.55 Δ = 0.15
R = 0.91 β = 0.35 Δ = 0.12
R = 0.86 β = 0.21 Δ = 0.06
R = 0.93 β = 0.42 Δ = 0.06
R = 0.92 β = 0.46 Δ = 0.06
R = 0.9 β = 0.25 Δ = 0.06
R = 0.86 β = 0.15 Δ = 0.09
R = 0.88 β = 0.48 Δ = 0.52
R = 0.89 β = 0.42 Δ = 0.04
R = 0.87 β = 0.15 Δ = 0.14
R = 0.81 β = 0.03 Δ = 0.18
R = 0.91 β = 0.78 Δ = 0.04
R = 0.91 β = 0.67 Δ = 0.03
R = 0.85 β = 0.16 Δ = 0.16
R = 0.85 β = 0.07 Δ = 0.18
R = 0.89 β = 0.76 Δ = 0.03
R = 0.87 β = 0.55 Δ = 0.08
R = 0.82 β = 0.1 Δ = 0.17
R = 0.81 β = 0.04 Δ = 0.19

**Figure S2. Full genome ancestry estimated by method for discovery dataset (N = 176).** Correlations between global ancestry proportions estimated by ADMIXTURE and LAI methods vs qpAdm. Each point represents one of the 176 admixed Neolithic individuals. R denotes the Pearson correlation between the method and qpAdm individual estimates. β indicates the slope of the regression line between the method and qpAdm estimates. Δ represents the average absolute difference in global ancestry between the LAI method and qpAdm. The dashed line corresponds to y = x for qpAdm estimates, while the red line shows the best-fit linear regression between qpAdm and the LAI method. Sample sizes of sources are listed for each column in the format of 7 farmers v N hunter-gatherers (7vN). Rows are labeled by method and whether calls were filtered by 0.9 posterior probability.

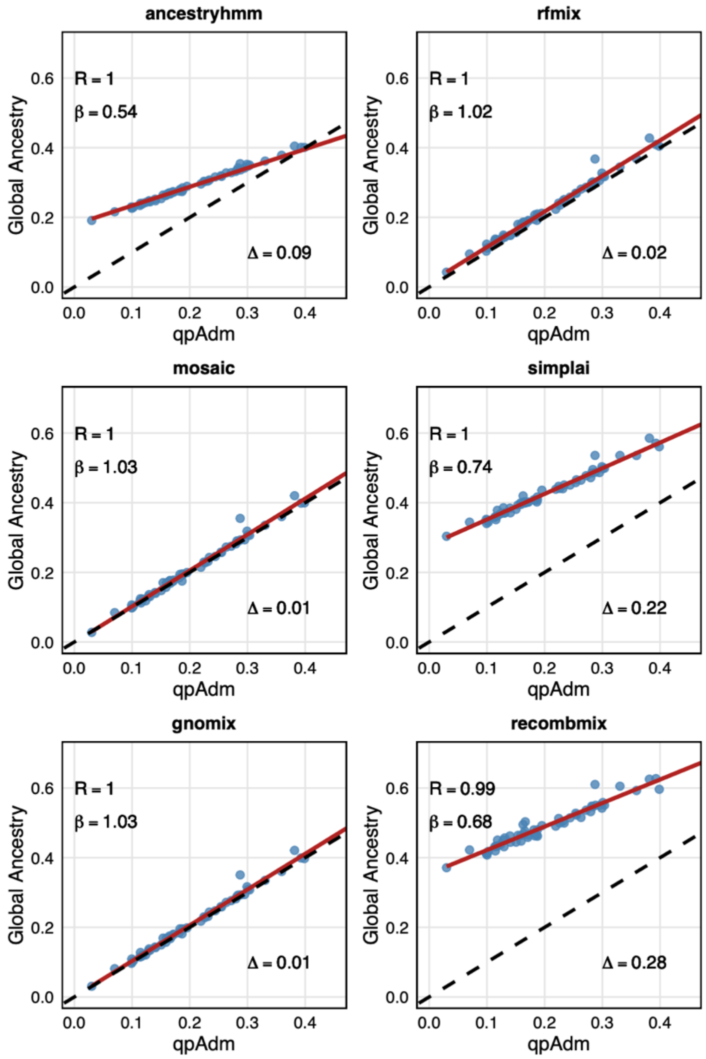


**Figure S3. Full genome ancestry estimated by LAI methods in 1000 Genomes Project African Americans.** Correlations in European (CEU) global ancestry proportions between LAI method and qpAdm. Each point is one of the 61 admixed African American (ASW) individuals.

R denotes the Pearson correlation between the method and qpAdm individual estimates. β indicates the slope of the regression line between the method and qpAdm estimates. Δ represents the average absolute difference in global ancestry between the LAI method and qpAdm. The dashed line corresponds to y = x for qpAdm estimates, while the red line shows the best-fit linear regression between qpAdm and the LAI method. Sample sizes of sources are 7 European (CEU) and 48 African (YRI) sources for Ancestry HMM, RFMix, Mosaic, Gnomix, Recomb-Mix, and 7v7 for simpLAI.

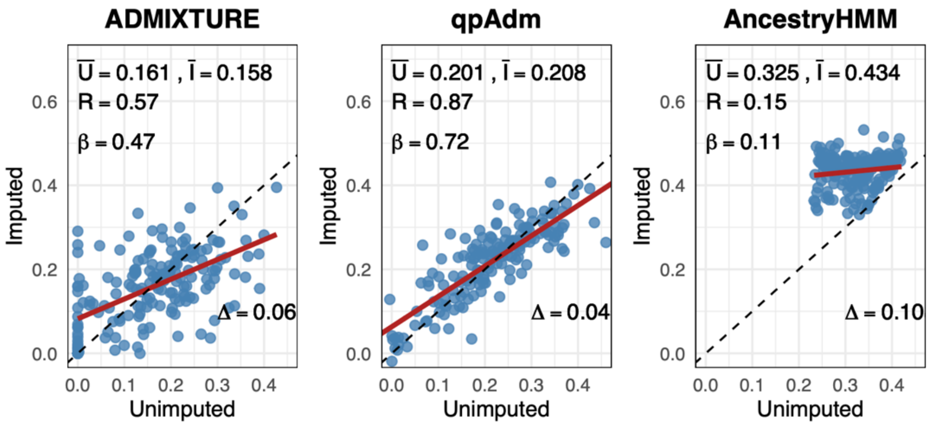


**Figure S4. Full genome ancestry estimated for imputed and unimputed data**. Figures show correlations in hunter-gatherer global ancestry proportions comparing results from **A)** qpAdm, **B)** ADMIXTURE, and **C)** Ancestry HMM on imputed and unimputed data. Each point is one of the 176 admixed Neolithic individuals. Ū and Ī estimates are mean ancestry estimates for unimputed and imputed data for each method respectively. R denotes the Pearson correlation between the method's individual estimates for unimputed and imputed datatypes. β indicates the slope of the regression line between the method's two datatypes individual estimates. Δ represents the average absolute difference in global ancestry between the method on each datatype. The dashed line corresponds to y = x for unimputed datatype results, while the red line shows the best-fit linear regression between method on unimputed and imputed data. Sample sizes of sources are 7 farmers and 48 hunter-gatherers.

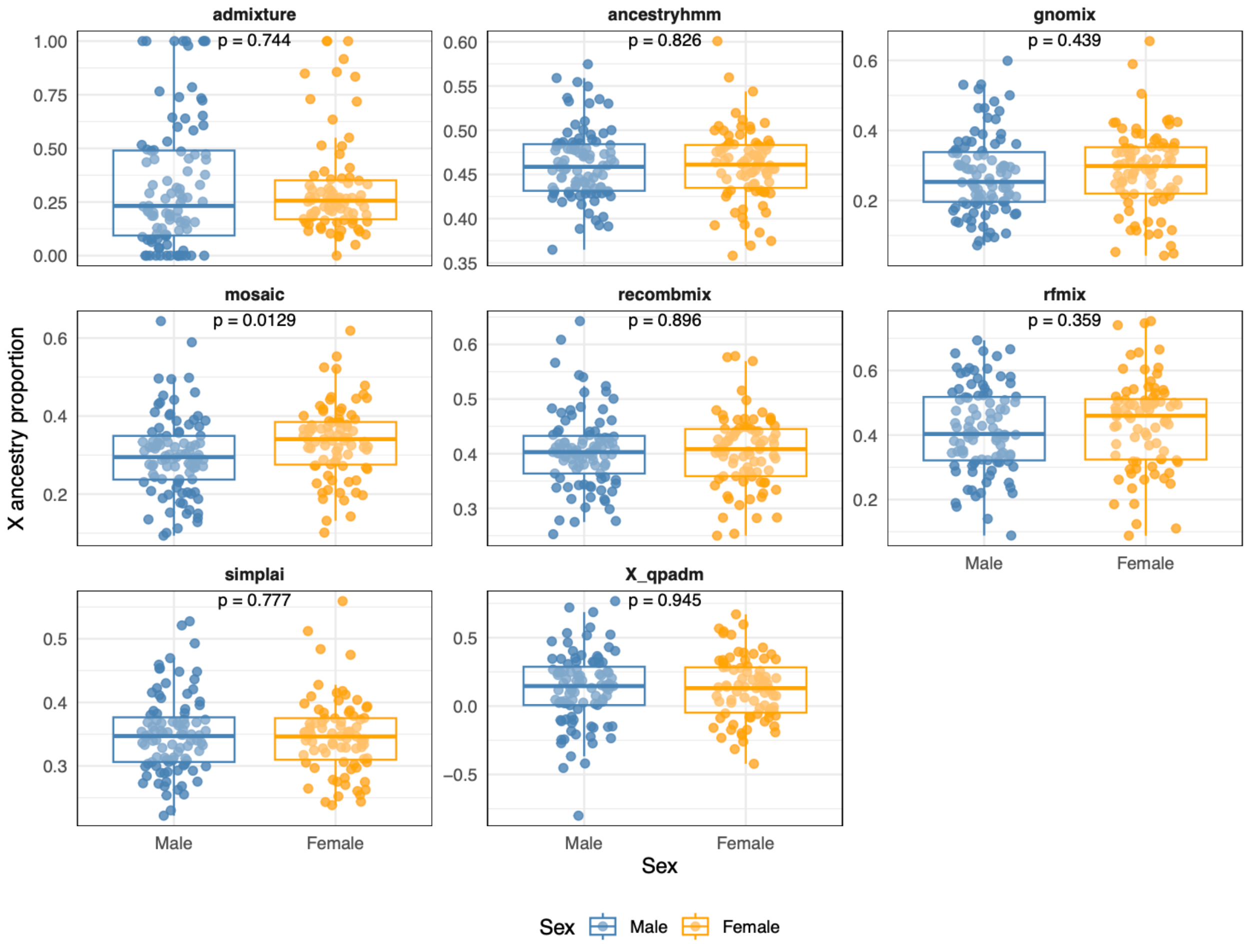


**Figure S5. X chromosome discovery dataset (N = 176) hunter-gatherer ancestry by sex.** Genome-wide X chromosome ancestry proportions for males and females across LAI methods for discovery dataset. Source sample sizes follow 7vN used in Figure 2. P-values are from two-sided t-tests comparing mean ancestry between sexes. Boxplots show median and interquartile range.

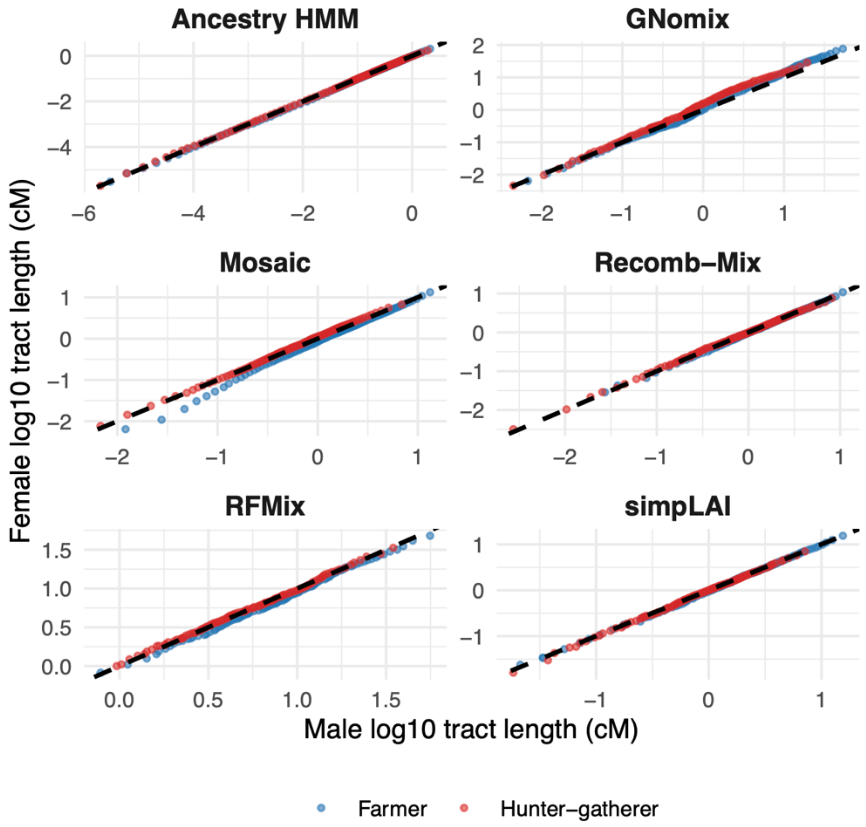


**Figure S6. QQ plot of X chromosome tract length distribution for discovery dataset (N = 176).** Comparing male and female tract length distributions (in log10 tract length (cM)) for each method, shown separately for farmer and hunter-gatherer ancestries.

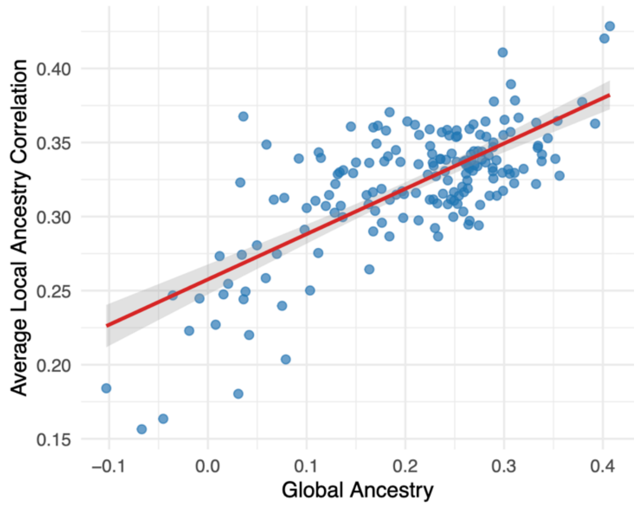


**Figure S7. Individual-level average local ancestry correlation across method pairs in discovery dataset (N = 176).** Individual-level average pairwise local ancestry correlation across seven methods, computed in 100 kb bins and plotted against global hunter-gatherer ancestry (qpAdm). Each point represents an individual. The red line indicates the best-fit linear regression. Source sample sizes follow 7vN used in Figure 2.

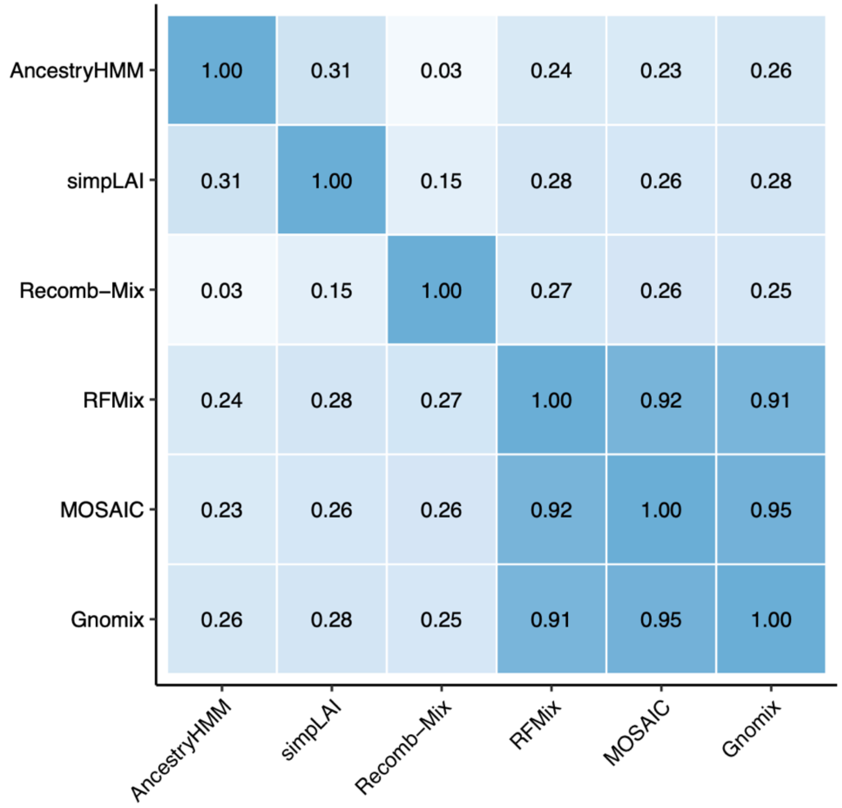


**Figure S8. Local ancestry correlations for LAI methods in 1000 Genomes.** Correlations in European (CEU) local ancestry proportions between LAI methods in 100kb bins along the genome. Sample sizes of sources are 7 CEU and 48 YRI sources for Ancestry HMM, RFMix, Gnomix, Recomb-Mix and Mosaic, and 7v7 for simpLAI.

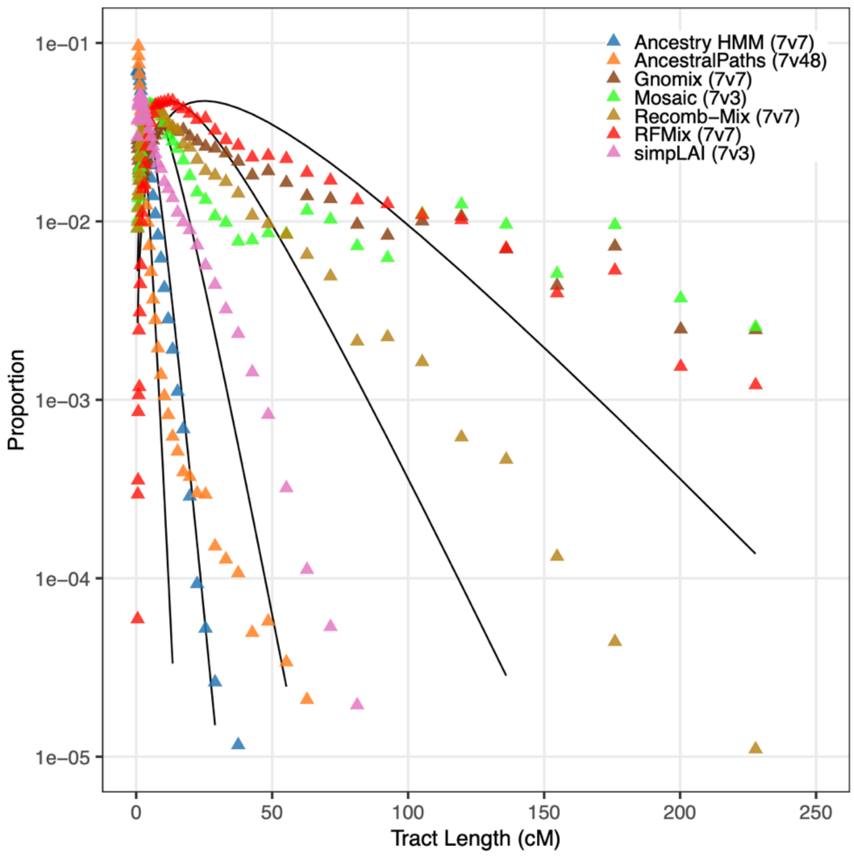


**Figure S9. Farmer tract length distributions by local ancestry inference method in discovery dataset (N = 176).** Proportion of farmer ancestry tracts by length (in cM) inferred using seven local ancestry inference methods. Empirical tract length distributions are shown as points, and black lines indicate theoretical single-pulse admixture models with admixture times (T) of 200, 100, 50, 20, and 10 generations. The figure provides the corresponding farmer ancestry tract distributions to complement the hunter-gatherer tract length analysis presented in Figure 4 of the main text. Source sample sizes are indicated in parentheses with each method title, representing 7 farmer reference individuals versus the number hunter gatherer sources (7vN).

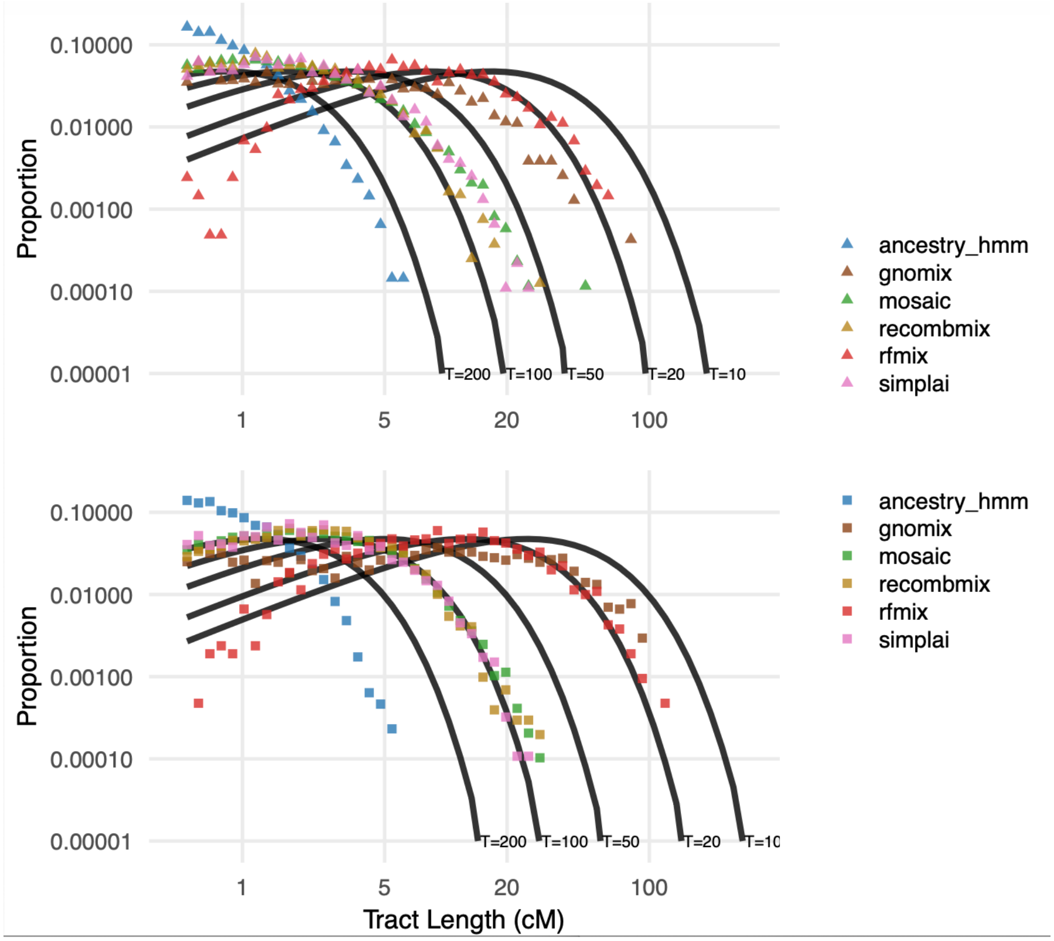


**Figure S10. X-chromosome tract length distributions inferred by LAI methods for discovery dataset (N = 176).** Distribution of tract lengths (cM) for each method compared to theoretical expectations (T = 200, 100, 50, 20, 10 generations). Top panel shows hunter-gatherer ancestry tract lengths (triangles) and bottom panel shows farmer ancestry tract lengths (squares).

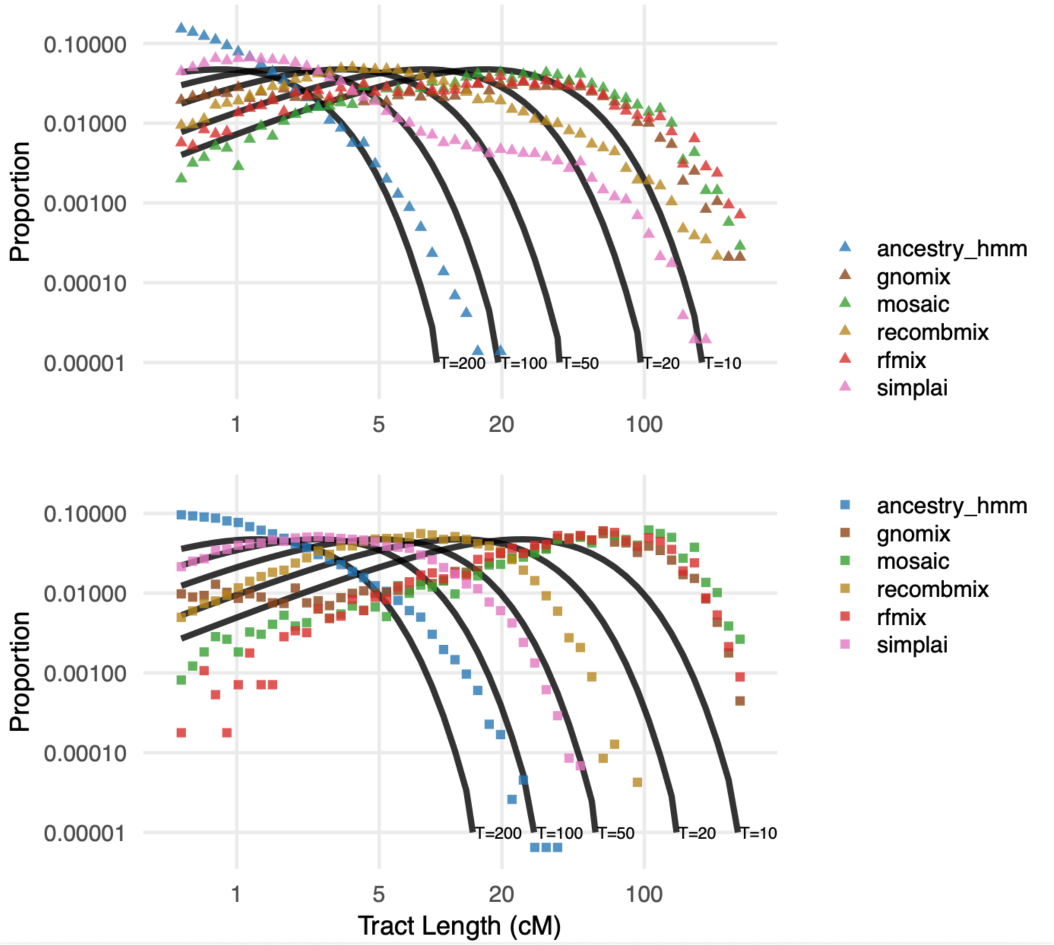


**Figure S11. Tract length distributions inferred by LAI methods in 1000 Genomes Project African Americans.** Distribution of tract lengths (cM) for each method compared to theoretical expectations (T = 200, 100, 50, 20, 10 generations). Top panel shows CEU ancestry tract lengths (triangles) and bottom panel shows YRI ancestry tract lengths (squares). Sample sizes of sources are 7 CEU and 48 YRI sources for Ancestry HMM, RFMix, Gnomix, Recomb-Mix, and Mosaic, and 7v7 for simpLAI.

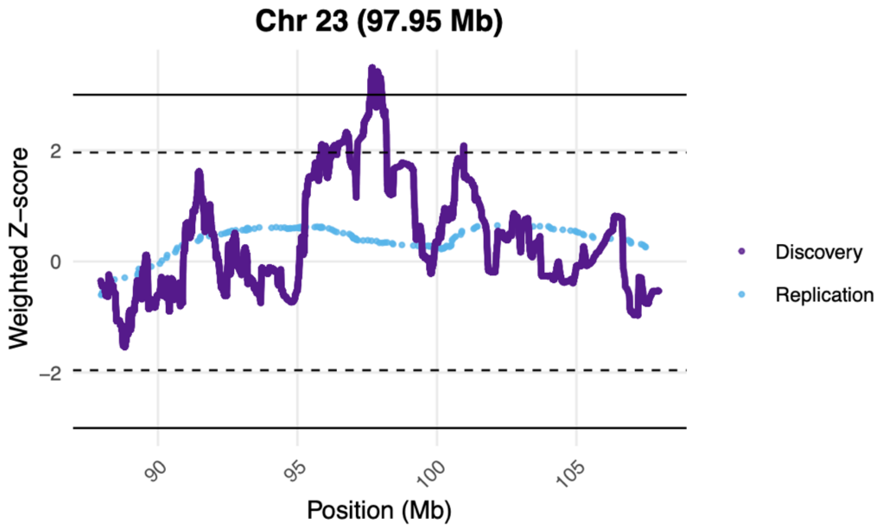


**Figure S12. Replication of X chromosome hit.** Comparison of combined and replication Z-scores for X chromosome loci exceeding the Z = ±3 discovery threshold. Solid lines indicate the discovery threshold (Z = ±3), and dashed lines indicate the replication threshold (Z = ±1.96). Discovery results (dark purple) are from meta-analysis of the six local ancestry inference methods on X chromosome discovery analyses (N = 176; Mosaic, RFMix, Gnomix, Recomb-Mix, simpLAI), and replication results (light blue) are from unimputed replication dataset 1 analyzed with Ancestry HMM.

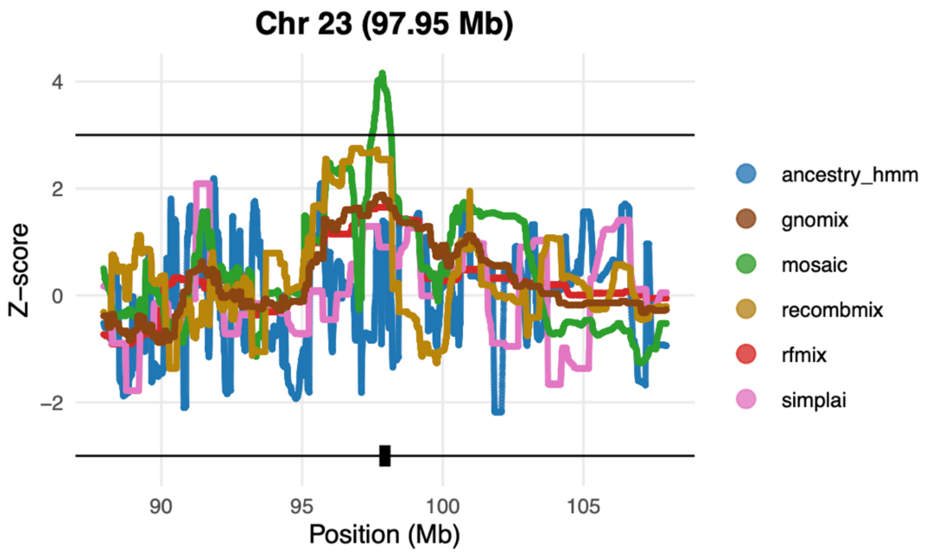


**Figure S13. Z-score deviations across methods for X chromosome hit.** Z-scores across methods for the discovery dataset (N = 176) X chromosome region (X:97.95 Mb). Horizontal lines indicate Z = ±3 thresholds used in the meta-analysis.

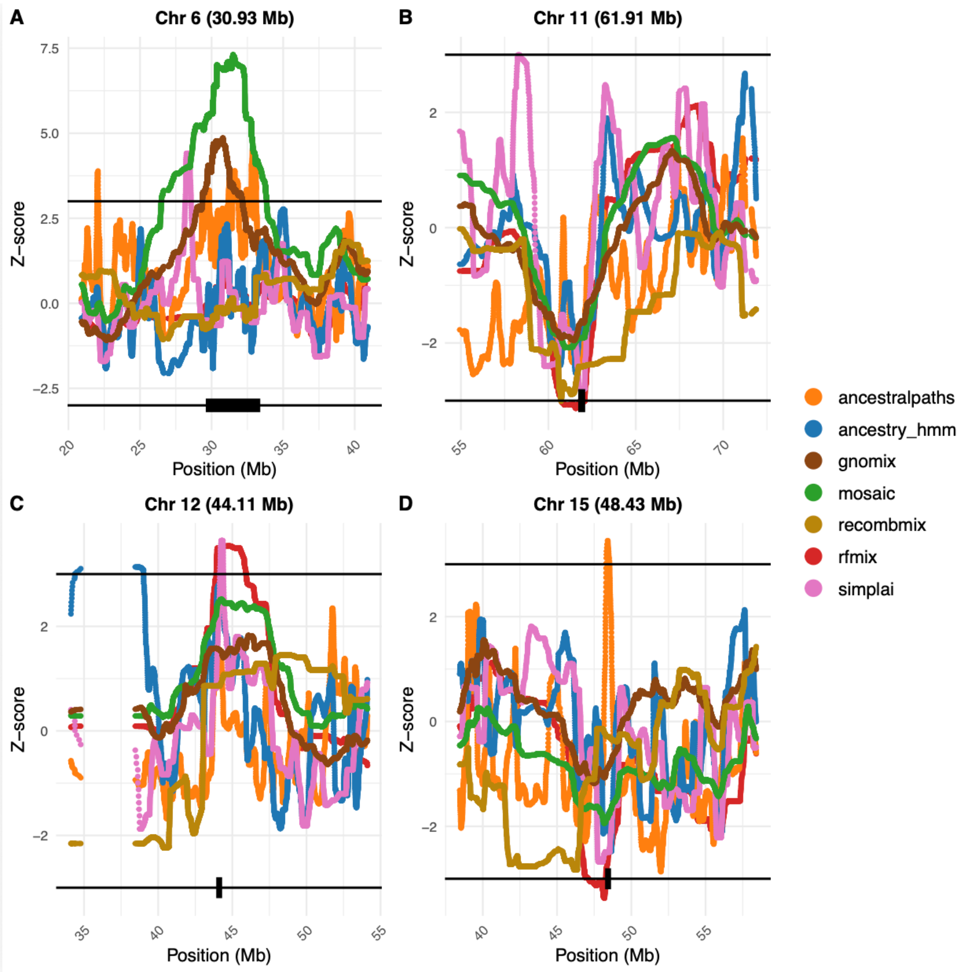


**Figure S14. Z-score deviations across methods for replicated loci.** Z-scores across methods for the five replicated regions in the discovery dataset. Horizontal lines indicate Z = ±3 thresholds used in the meta-analysis. Methods use 7v48 source panels, except simpLAI (7v7). Putative candidate genes are highlighted with black boxes.

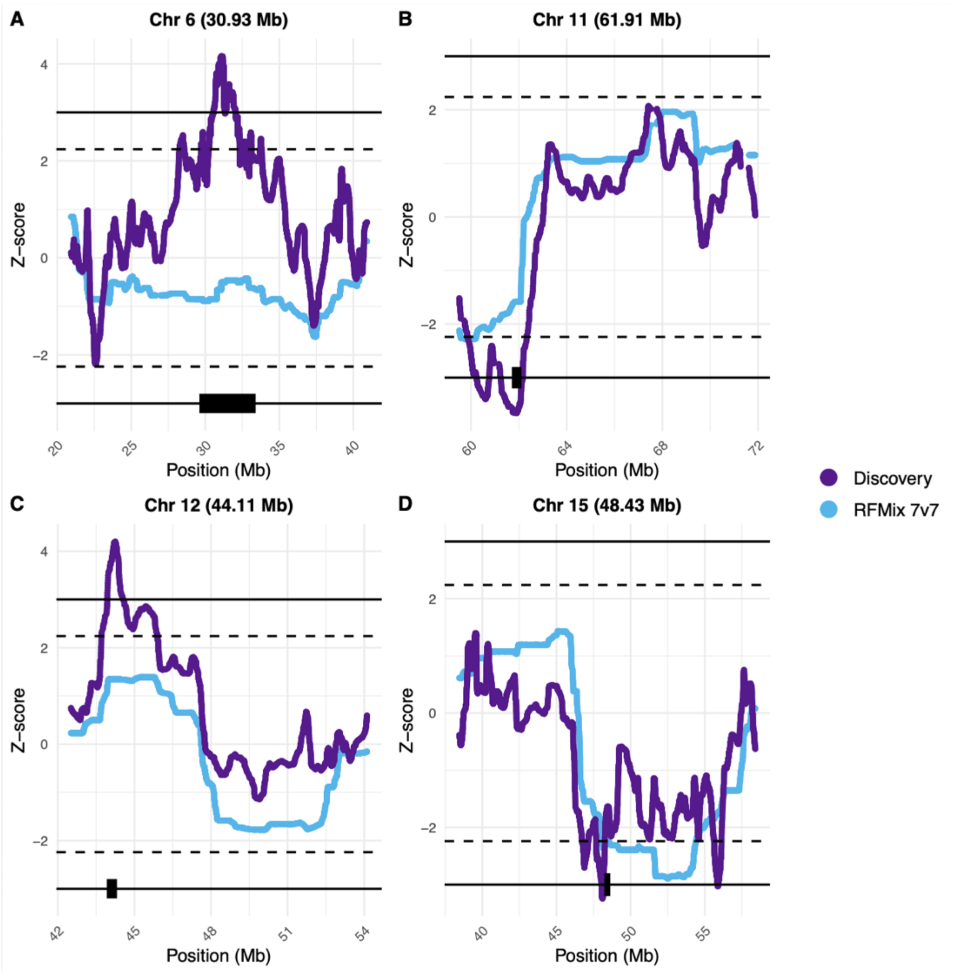


**Figure S15. Replication of Z-scores from meta-analysis of discovery dataset in RFMix 7v7 results.** Comparison of meta-analysis of seven methods with 7v48 sources Z-scores and RFMix 7v7 Z-scores at the four replicated loci. Dashed lines indicate the discovery threshold ($Z = \pm 3$), and solid lines indicate the replication threshold ($Z = \pm 2.64$).

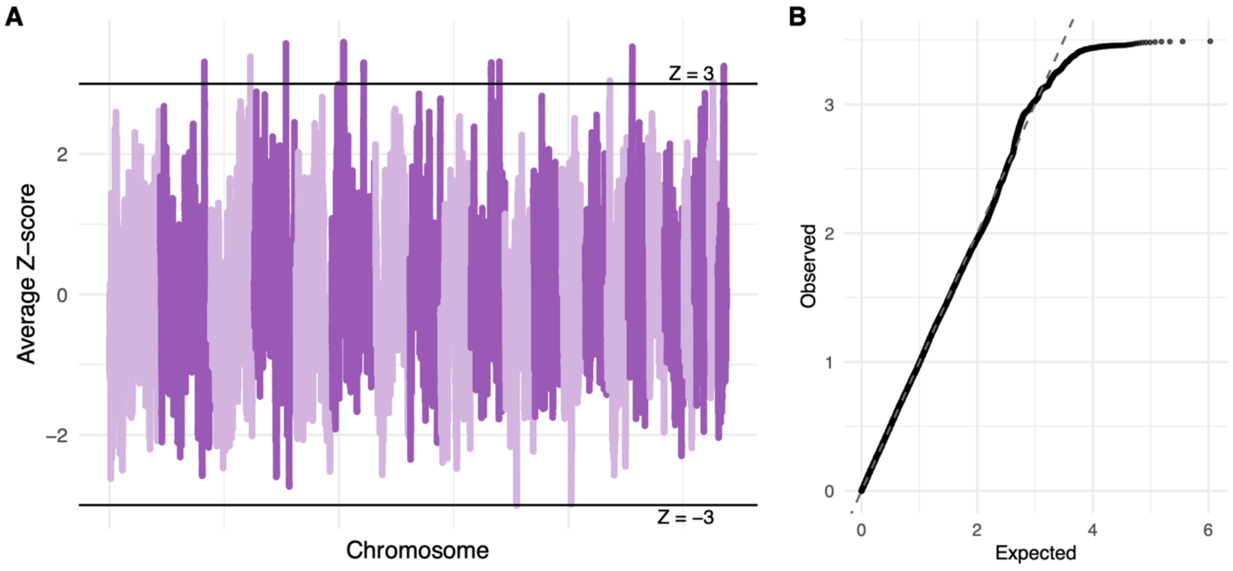


**Figure S16. Manhattan and QQ plots for meta-analysis of discovery dataset (N = 176) using 7vN source reference panels.** Results for all methods using 7vN source panels. Horizontal dashed lines indicate Z = ±3. Source sample sizes (7vN) correspond to Figure 2.

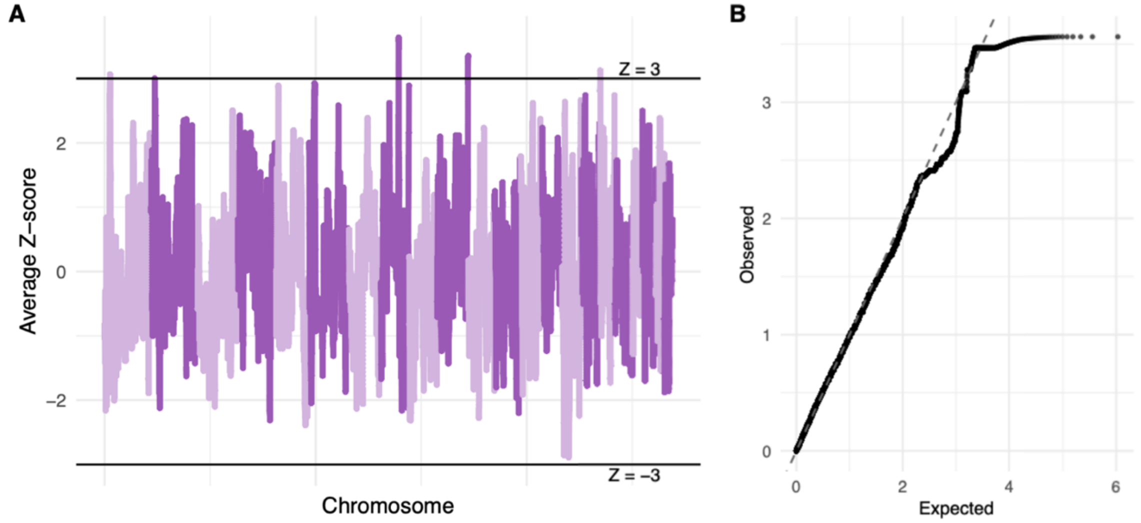


**Figure S17. Manhattan and QQ plots for RFMix 7v7.** Horizontal dashed lines indicate Z = ±3.
